# Tracing back the expansion of the endomembrane rescue systems in parasitic Parabasalids to the ancestral origin in its free-living sister lineage

**DOI:** 10.1101/2024.01.09.574884

**Authors:** Abhishek Prakash Shinde, Jitka Kučerová, Joel Bryan Dacks, Jan Tachezy

**Affiliations:** Department of Parasitology, Faculty of Science, Charles University, BIOCEV, Průmyslová 595, 25242 Vestec, Czech Republic; Division of Infectious Diseases, Department of Medicine and Department of Biological Sciences, University of Alberta, Edmonton, Alberta, Canada, University of Alberta, Edmonton, Canada; Women and Children’s Health Research Institute, University of Alberta; Department of Genetics, Evolution, & Environment, University College, London; Institute of Parasitology, Biology Centre, Czech Academy of Sciences, České Budějovice (Budweis), Czech Republic

**Keywords:** Retromer, Retriever, Phylogenomics, Endomembrane, Evolution, Parabasalids

## Abstract

Early Endosomes sort transmembrane cargo whether for lysosomal degradation or retrieval to the plasma membrane or the Golgi complex. Endosomal retrieval in eukaryotes is governed by the anciently homologous Retromer or Retriever complexes. Each comprises a core tri-protein subcomplex, membrane-deformation proteins, and interacting partner complexes, together retrieving a variety of known cargo proteins. *Trichomonas vaginalis*; a sexually transmitted human parasite uses the endomembrane system for pathogenesis. It has massively and selectively expanded its endomembrane protein complement, the evolutionary path of which has been largely unexplored. Our molecular evolutionary study of Retromer, Retriever and associated machinery in parabasalids and its free-living sister lineage of Anaeramoeba, demonstrates specific expansion of the Retromer machinery, contrasting with the Retriever components. We also observe partial loss of Commander complex and Sorting Nexins in Parabasalia but complete retention in Anaeramoeba. Notably, we identify putative parabasalid Sorting Nexin analogues. Finally, we report the first Retriever protein localization in a non-metazoan group along with Retromer protein localization in *T. vaginalis*. Therefore, we provide a unique genome expansion study of endomembrane trafficking system in parasitic and free-living protists alike.

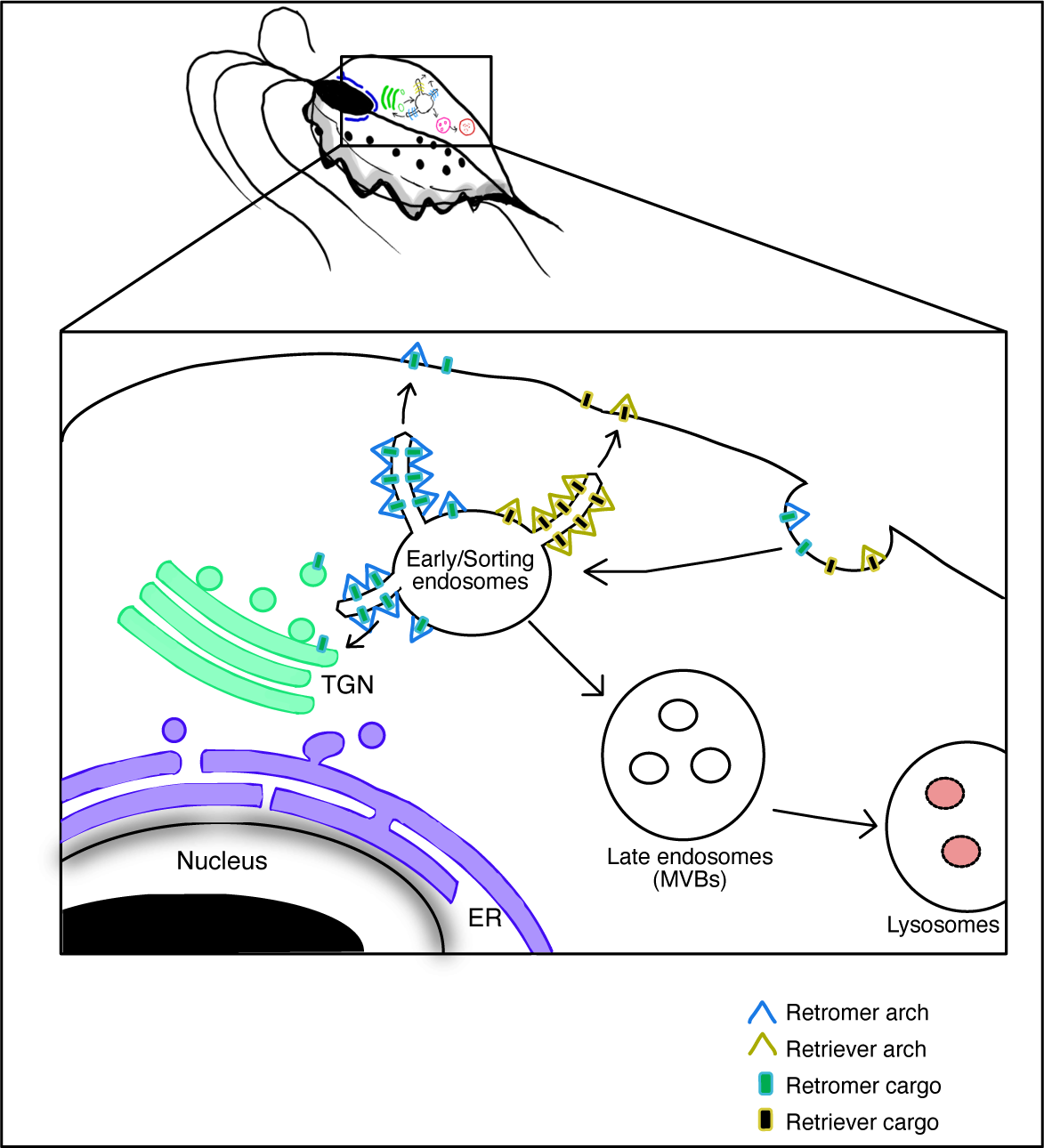

## 1. INTRODUCTION

*Trichomonas vaginalis* is a parasitic flagellated protist that causes the commonly diagnosed non-viral sexually transmitted infection generally known as trichomoniasis. In the year 2020, 156 million new cases of this infection among people aged between 15 to 49 years were registered worldwide as surveyed by the World Health Organization (WHO). In females, a variety of symptoms are observed, such as vaginitis and cervicitis. Along with these symptoms, adverse outcomes include complications in pregnancy such as preterm birth, low birth weight, cervical cancer, and a 1.5-fold higher risk of contracting an HIV infection (Edwards et al., 2016), (McClelland et al., 2007). It is also reported that trichomoniasis infection during pregnancy can lead to intellectual disability in children (Mann et al., 2009). Though generally asymptomatic in men, it can cause urethral inflammation and increase the risk of prostate cancer (Sutcliffe et al., 2006), (Krieger, 1995).

*T. vaginalis* is an anaerobic parasitic protist belonging to the monophyletic Metamonada group and Parabasalia subgroup (Burki et al., 2020), (Adl et al., 2019). Parabasalia are flagellated organisms distinguished by the presence of hydrogenosomes, an anaerobic type of mitochondria, a cytoskeletal framework of unique microtubular elements known as the pelta-axostylar system, and a parabasal apparatus (Céza et al., 2022), (Benchimol et al., 2000). Two distinct Parabasalia lineages; Trichomonadea and Tritrichomonadea have majorly adopted a parasitic lifestyle (Malik et al., 2011). Recently, free-living Anaeramoebae were shown to represent the closest sister lineage of Parabasalia and found to serve a good contrast of lifestyles between related organisms (Stairs et al., 2021), (Maciejowski et al., 2023).

*T. vaginalis* exhibits a massive genome expansion of selective gene families, including the Membrane Trafficking System (MTS) involved in endocytosis and exocytosis of the cell, it also shows expansion of several of its secreted virulence factors (Jane M. Carlton & Robert P. Hirt, n.d.). The endomembrane machinery of *T. vaginalis* plays a crucial role in its pathogenesis. For instance, the lysosomal secretion of its virulence factors such as cysteine peptidases and Cathepsin D is guided by the endosomal MTS of the cell (Zimmann et al., 2022). To maintain this continuous secretion of virulence factors, endomembrane proteins responsible for their anterograde transport are expected to be rescued by retrograde trafficking to avoid their lysosomal degradation.

In eukaryotes, the MTS constitutes anterograde and retrograde trafficking; while anterograde trafficking transports cargoes from the endoplasmic reticulum (ER) through the trans-Golgi network (TGN) to the plasma membrane, retrograde trafficking recycles the cargoes by sorting them in the early endosomes. This sorting of cargo in the endosomes helps with the recovery of essential resident endomembrane proteins required by the cell and are used for new rounds of trafficking while discarding the foreign material via lysosomal degradation through the endosomal sorting complexes required for transport (ESCRT) machinery (Raiborg & Stenmark, 2009),(Pipaliya et al., 2021), (Huotari & Helenius, 2011).

Endosomal retrograde trafficking was first reported to be carried out by a membrane coat complex, which was identified and characterized for the rescue of Vacuolar protein sorting receptor Vps10p, a sortilin homolog in yeast (Seaman et al., 1998). Vacuolar protein sorting proteins Vps35p and Vps29p were shown to form a multimeric complex with Vps26p, Vps17p, and Vps5p, which was collectively termed the Retromer complex. The hetero-trimeric complex of Vps35p, Vps29p, and Vps26p conducts cargo selection for retrieval, while Vps5p and Vps17p assembly promotes vesicle formation via endomembrane deformation (Seaman et al., 1998). This was followed by the discovery of VPS29, VPS35 homologs and two VPS26A/B homologs in humans (Edgar & Polak, 2000), (HaP et al., 2000), (Kerr et al., 2005). Indeed, the hetero-trimeric subcomplex proteins of the Retromer complex were confirmed to be highly conserved across pan-eukaryotic species (Koumandou et al., 2011).

In mammalian systems, the term Retromer complex means only the conserved trimeric complex of VPS26, VPS29, and VPS35 (Seaman, 2021). In humans, the Retromer complex was primarily shown to rescue Cation-independent Mannose 6-phosphate receptor (CIMPR) responsible for transporting lysosomal hydrolases possessing Mannose 6-phosphate signal from TGN to late endosomes (Seaman et al., 1998), (Edgar & Polak, 2000), (Arighi et al., 2004). Other than CIMPR and Vps10p, Retromer complex also rescues late-Golgi enzyme Kexin2 (KEX2) and a Golgi-resident endopeptidase, dipeptidyl amino peptidase (DPAP) (Seaman et al., 1998).

For a long time, the Retromer complex was believed to be the only endosomal retrograde trafficking machinery. That changed with the recent discovery of the Retriever complex in 2017; Retromer-independent rescue machinery in the mammalian system (McNally et al., 2017). The Retriever complex is also a hetero-trimeric complex, structurally homologous to the Retromer core complex. It consists of DSCR3, C16orf62 or VPS35-like (VPS35-L) and the shared VPS29 (McNally et al., 2017), (Chen et al., 2019). It only recycles plasma membrane proteins missed by the Retromer complex in mammalian cells, such as low-density lipoprotein receptors (LDLR), LDLR-like protein (LRP-1), and χξ5ϕ31 integrin (McNally et al., 2017), (McNally & Cullen, 2018).

In addition to the core trimeric Retromer and Retriever complexes, the rescue of endomembrane proteins also depends on Sorting Nexins or Cargo adaptors, they recruit the trimeric complexes to associate with the endomembrane cargo. They also mediate vesicle formation via endomembrane deformation. First identified in yeast assisting the Retromer complex; were hetero-dimeric Vps5p and Vps17p, both of which possess a PX (Phox homology) domain and a BAR (Bin, Amphiphysin, and Rvs) domain (Nothwehr & Hindes, 1997), (Teasdale & Collins, 2012), (Lucas et al., 2016). In mammals, all the Sorting Nexins besides possessing a lipid binding PX domain, based on the presence or absence of additional domains are categorized as SNX-PX, SNX-BAR, and SNX-FERM. All three categories of Sorting Nexins are known to assist the Retromer complex. Only SNX-FERM is known to assist the Retriever complex (Harrison et al., 2014), (Steinberg et al., 2013), (Strochlic et al., 2007). However, no SNX proteins were previously identified in *T. vaginalis* (Koumandou et al., 2011).

Among the accessory factors shared by both the rescue machineries, an essential one is the pentameric <u>W</u>iskott-<u>A</u>ldrich syndrome protein and <u>S</u>CAR <u>H</u>omologue (WASH) complex which is responsible for actin polymerization along the endosomal membrane to mediate endomembrane tubulation (Derivery et al., 2009), (Healy et al., 2023). Apart from the WASH complex, Retriever machinery also requires another essential accessory factor, a huge complex of 13 proteins known as the CCC complex comprising of CCDC22, CCDC93, 10 COMMD (<u>Co</u>pper <u>M</u>etabolism <u>M</u>URR1 [Mouse U2af1-rs1 region 1] <u>D</u>omain) proteins and DENND10 (<u>D</u>ifferentially <u>E</u>xpressed in <u>N</u>ormal and <u>N</u>eoplastic cells-containing <u>D</u>omain protein 10 / FAM45A). Seeing as how Retriever and CCC complexes interact and co-depend on each other, both these complexes have been termed together as the Commander complex (Mallam & Marcotte, 2017), (Healy et al., 2023).

Given the expansion of the Retromer complex, the reported presence of the Retriever component VPS35L in *T. vaginalis*, and the new availability of parabasalid genomes since the last time that these complexes were examined, we decided to undertake a molecular evolutionary analysis to determine the path that has produced this unusual protein complement. Moreover, since the endosomal system is involved in *T. vaginalis* pathogenesis, and since the Retriever complex has never been characterized outside of mammalian systems, we investigated the relative localization of Retromer vs Retriever components to determine whether there are multiple endosomal retrieval pathways.

## 2. RESULTS

### 2.1 Retromer complex genes are widely expanded in Parabasalids and Anaeramoeba compared to the Retriever complex

In the previous studies conducted for the genomic survey of Retromer complex across pan-eukaryotic species, *T. vaginalis* was the only Metamonad that showed multiple gene paralogues of trimeric Retromer complex proteins; VPS26A/B, VPS29, and VPS35 (Koumandou et al., 2011). Pan-eukaryotic distribution of the Retriever complex was limited only to VPS35L (McNally et al., 2017). So, we decided to perform a deep dive genomic analysis into the expansion of the Retromer complex, meanwhile searching for the Retriever complex and both their specific endomembrane cargoes in parasitic parabasalids and their closest free-living sister *Anaeramoeba ignava* (Figure 1A).

**Figure 1:**
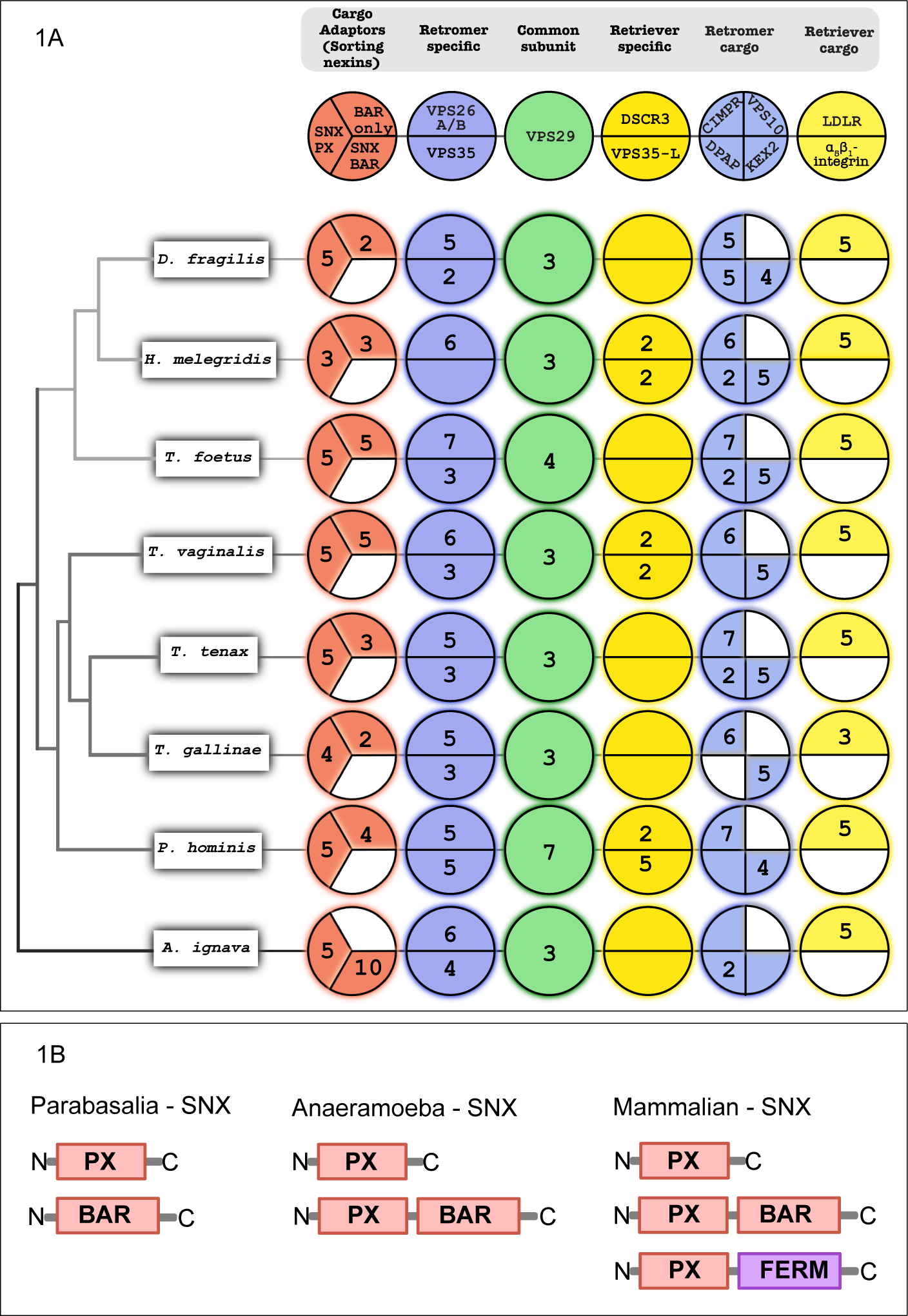
A) Coulson plot representing comparative genomics of the endosomal Retrograde trafficking proteins and their specific known cargoes across selected Parabasalia organisms and *Anaeramoeba ignava*. Areas filled with color demonstrate positive occurrence of the protein homologues, numbers indicate the paralogue count and colored sections with no numbers indicate a single paralogue. Sections left white or colorless indicate the absence of protein homologues. Each plot column is annotated with protein name at the top and the name of organism is mentioned on the left. B) Comparison of sorting nexins (SNX) identified in these organisms with the canonical mammalian SNX; Parabasalia contain SNX-PX and a unique SNX; homologous to VPS5 C-terminus BAR domain. Anaeramoeba contain SNX-PX and SNX-BAR, while Humans have an additional SNX-FERM.

In all the Parabasalia members, we observed an expansion of VPS26A/B ranging from five genes in *Trichomonas gallinae* and *Dientamoeba fragilis* to seven genes in *Tritrichomonas foetus*. Similarly, for VPS35, we observed the expansion ranging from 2-5 genes in nearly all parabasalids. The shared protein between Retromer and Retriever complexes; VPS29, was seen to be expanded ranging from 3-4 genes in all Parabasalia members, except for seven genes in *Pentatrichomonas hominis.* Somewhat unexpectedly, the gene expansions of Retromer complex proteins were found also in the free-living *A. ignava* with six and four genes of VPS26A/B and VPS35 respectively, along with three genes for VPS29 (Figure 1A).

A bigger surprise for us was seeing almost a complete lack of gene expansions for Retriever complex-specific proteins; DSCR3 and VPS35-L. In nearly all parabasalids and *A. ignava*, only a single gene was found for both proteins. However, two copies were found in *H. meleagridis* and *T. vaginalis*, except for *P. hominis*, in which besides the two copies of DSCR3 we observed five copies of VPS35-L.

Since DSCR3 and VPS35-L are ancient homologs of Retromer complex proteins; VPS26A/B and VPS35 respectively, are believed to be a result of ancestral duplication events in LECA (Koumandou et al., 2011), (McNally et al., 2017), we chose to validate our classifications via phylogenetics. This also allowed us to trace the origins of the observed expansions of the Retromer components in the parabasalids. We conducted phylogenetic analyses of the respective Retromer and Retriever complex homologs to investigate their classification in parabasalids and *A. ignava* (Figure 2).

**Figure 2:**
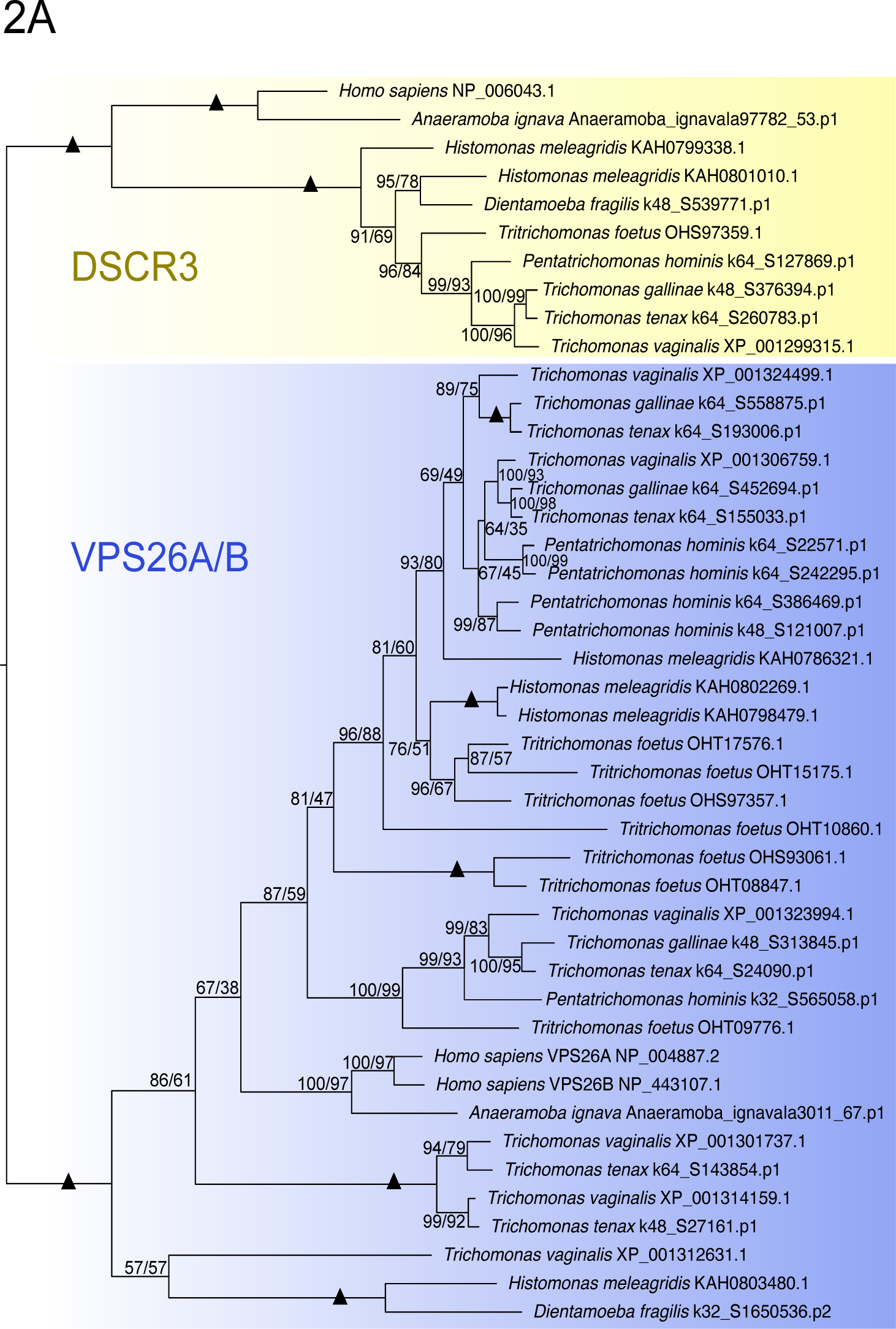
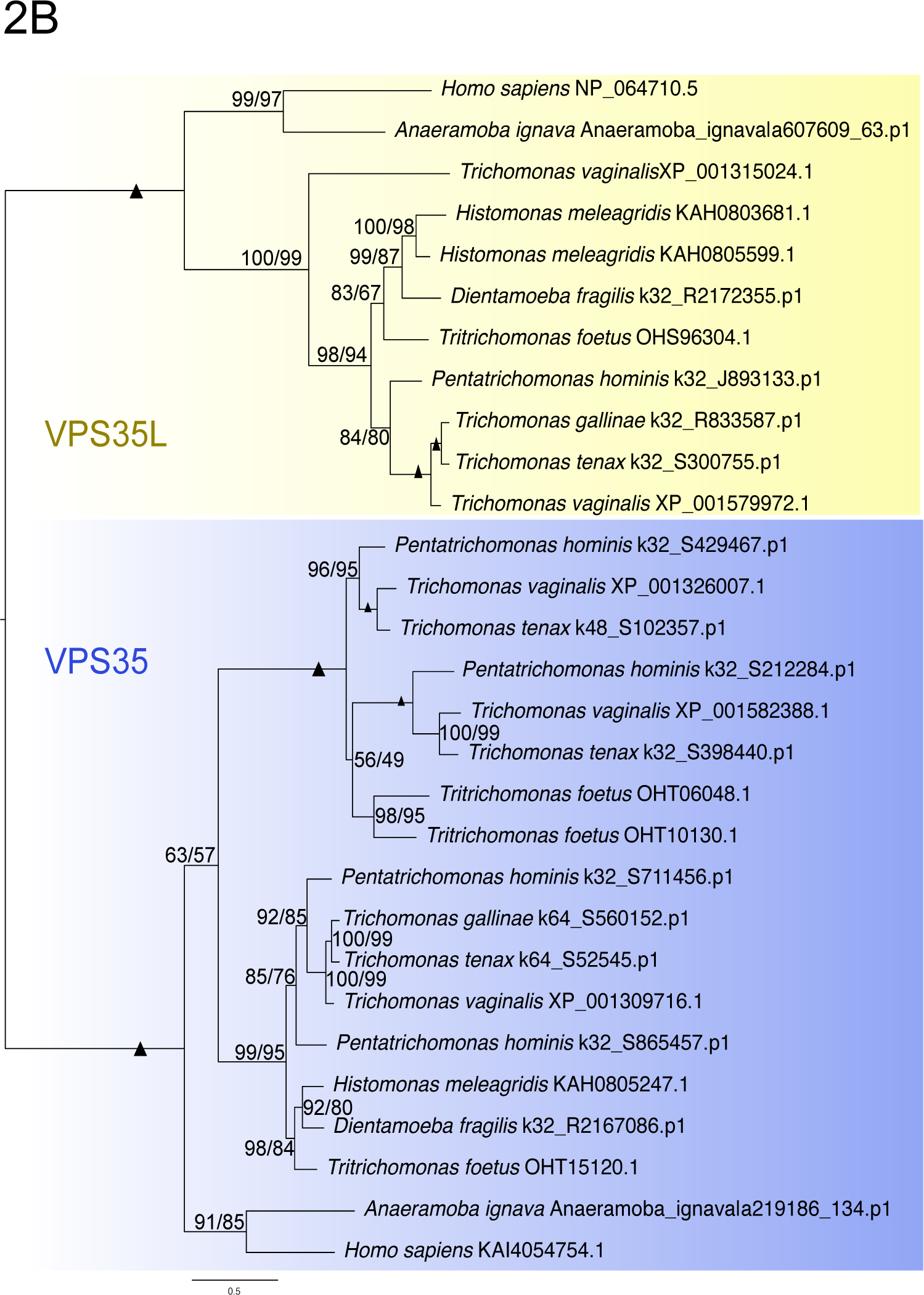
A) Maximum likelihood phylogenetic analysis tree was constructed using IQTree (Best fit: LG+F+G4) with 44 sequences and 222 sites for VPS26A/B and DSCR3. This tree includes all the orthologues of VPS26A/B and DSCR3 identified in the comparative genomics for Parabasalids. B) Maximum likelihood phylogenetic analysis tree was constructed using IQTree (Best fit: Q.yeast+G4) with 29 sequences and 487 sites for VPS35 and VPS35L. This tree includes all the orthologues of VPS35 and VPS35L identified in the comparative genomics for Parabasalids. *H. sapiens* and *A. ignava* sequences were used as a reference to the analyses. Clade in blue depicts the Retromer complex proteins and the clade in yellow depicts the Retriever complex proteins. Support values for each node are depicted as, UFB on the left indicates ultrafast bootstrap values and support values on the right indicate non-parametric (NP) bootstrap values (UFB/NP). Branches with support values of 100 for both ultrafast and NP bootstrapping are represented with black solid triangles.

We conducted phylogenetic characterization of VPS26A/B with DSCR3 (Figure 2A), and characterization of VPS35 with VPS35-L (Figure 2B), including the human paralogues as outgroups and functionally verified markers. This analysis shows separate clades of VPS26A/B and DSCR3 confirming the classification of our homology searches in parabasalids (Figure 2A). Similarly, VPS35 and VPS35-L are also seen to be classified in separate clades (Figure 2B). This also confirms that DSCR3 and VPS35-L of the Retriever complex have not been expanded. By contrast, VPS26 and VPS35 in parabasalids have been expanded via ancestral duplication events taking place prior to or in the Last Parabasalid Common Ancestor (LPCA).

We speculated from our homology searches that the expansion of the Retromer complex in parabasalids can be timed for the duplication events and possibly be traced back to *A. ignava*. Hence, we conducted phylogenetic analyses for each Retromer complex protein to trace their expansion in *A. ignava* and the parabasalids (Figure 3). For *A. ignava*, we carefully selected the sequences that showed the presence of conserved domains and ignored the partial sequences. All the *A. ignava* hits for VPS26A/B, showed the presence of a conserved domain, while only 2 hits of VPS29 and VPS35 each showed the complete presence of conserved domains. All the sequences from Parabasalid searches were included in the analyses.

**Figure 3:**
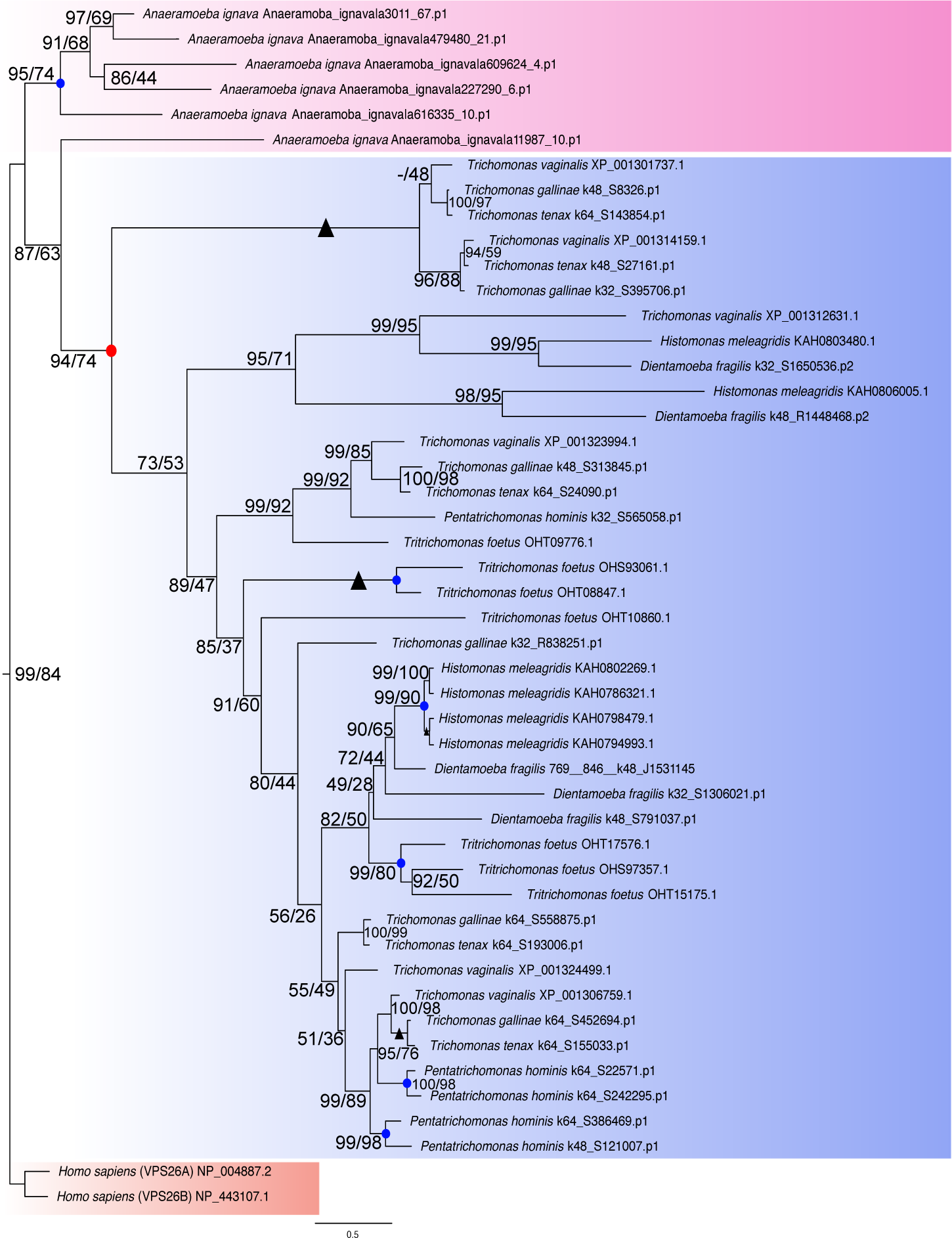
Maximum likelihood phylogenetic analysis tree was constructed using IQTree (Best fit: Q.yeast+I+G4) with 48 sequences and 270 sites for VPS26A/B expansion events in parasitic Parabasalia lineage and *A. ignava.* This includes all the orthologues identified from the homology searches. The clade in pink represents paralogues caused by species-specific expansion of VPS26A/B in *A. ignava*. The clade in blue represents orthologues of VPS26A/B identified in the parasitic Parabasalia lineage. VPS26A and VPS26B of *H. sapiens* were used as an outgroup represented by a red clade. The red dot represents an ancestral duplication event in parasitic Parabasalids while the blue dots represent species specific duplication events. Support values for each node are presented in the format of UFB/NP. Branches with support values of 100 for both ultrafast and NP bootstrapping are represented with black solid triangles.

In the phylogenetic analysis of VPS26A/B (Figure 3), we used VPS26A and VPS26B of humans as an outgroup. We observed that the expansion of VPS26A/B in *A. ignava* is independent of the expansion observed in Parabasalid members. Among parabasalids, the expansion of VPS26 appears to be a result of numerous ancestral duplication events. Also, species-specific duplication events appear to have occurred in *T. foetus*, *H. meleagridis*, and *P. hominis*. Expansion of the Retromer complex in *A. ignava* appears to be a result of either species-specific or lineage-specific duplication events.

After confirming the lineage-specific expansion of the Retromer complex in parabasalids and *A. ignava* for VPS26, we decided to use the *A. ignava* clade as an outgroup for the other analyses to increase robustness. We conducted phylogenetic analysis for the expansion of VPS29 and VPS35 of Retromer complex in parabasalids (Supporting Figures S1 & S2). We show an ancestral duplication event in LPCA resulting in two parabasalid specific homologs for both the proteins, i.e., VPS29A/B and VPS35A/B. Like the expansion of VPS26A/B, we observed species-specific duplication events in *P. hominis* and *T. foetus* for VPS29 and VPS35.

### 2.2 Parabasalids encode Retromer and Retriever complex-specific cargo proteins

As the Retromer complex proteins were seen to be expanded alongside contrasting observations for the Retriever complex specific proteins in Parabasalia members and *A. ignava*, we extended our searches to the specific endosomal membrane cargo proteins for both these machineries (Figure 1A). For the endomembrane cargoes of the Retromer complex, we looked for CIMPR, Vps10p (VPS10), DPAP, and KEX2. Consistent expansion was observed for CIMPR, with 5-7 genes in all the parasitic parabasalids, however, only a single gene of putative CIMPR was found in the free-living *A. ignava.* Similarly, KEX2 was also expanded with 4-5 genes in all parabasalids, while *A. ignava* has only a single gene. Expansion of DPAP was observed in all the Tritrichomonadea ranging from 2-5 genes, while a single gene was found in *T. vaginalis* and *P. hominis,* two genes in *T. tenax* and *A. ignava*, and completely absent in *T. gallinae*. VPS10 was seen to be completely missing in all taxa searched.

Next, we chose two functionally verified Retriever complex cargo candidates; LDLR and α_5_β_1_-integrin. While LDLR is known to be present in non-metazoan species, α_5_β_1_-integrin is thus far unique to humans, however, integrins have been identified within Amorphea (McNally et al., 2017), (Kang et al., 2021). Contrasting to the lack of expansion of Retriever machinery components, 3-5 genes of LDLR were found in all the parabasalids. However, LDLR was seen to be completely absent in *A. ignava*, but alternatively, five genes were found in *Anaeramoeba flamelloides* of the Anaeramoeba lineage. This could have been because of the fragmented datasets of *A. ignava* (Maciejowski et al., 2023). α_5_β_1_-integrin proteins were seen to be completely missing in both parabasalids and Anaeramoeba. These results suggest a similar pattern of evolution for both the rescue machineries in parasitic parabasalids and Anaeramoeba regardless of their contrasting lifestyles.

### 2.3 The evolution of the WASH complex and the CCC complex is consistent with the evolution of the Retriever complex in Parabasalia and Anaeramoeba lineages

The pentameric WASH complex responsible for actin polymerization along the endosomal membrane is known to be shared by Retromer and Retriever complexes and it plays a crucial role in the successful rescue of endomembrane cargoes by both these machineries. We conducted a comparative genomic survey of the WASH complex in Parabasalia and *A. ignava* genomes to understand the consistency of its evolutionary pattern with either of the hetero-trimeric complexes. (Figure 4).

**Figure 4:**
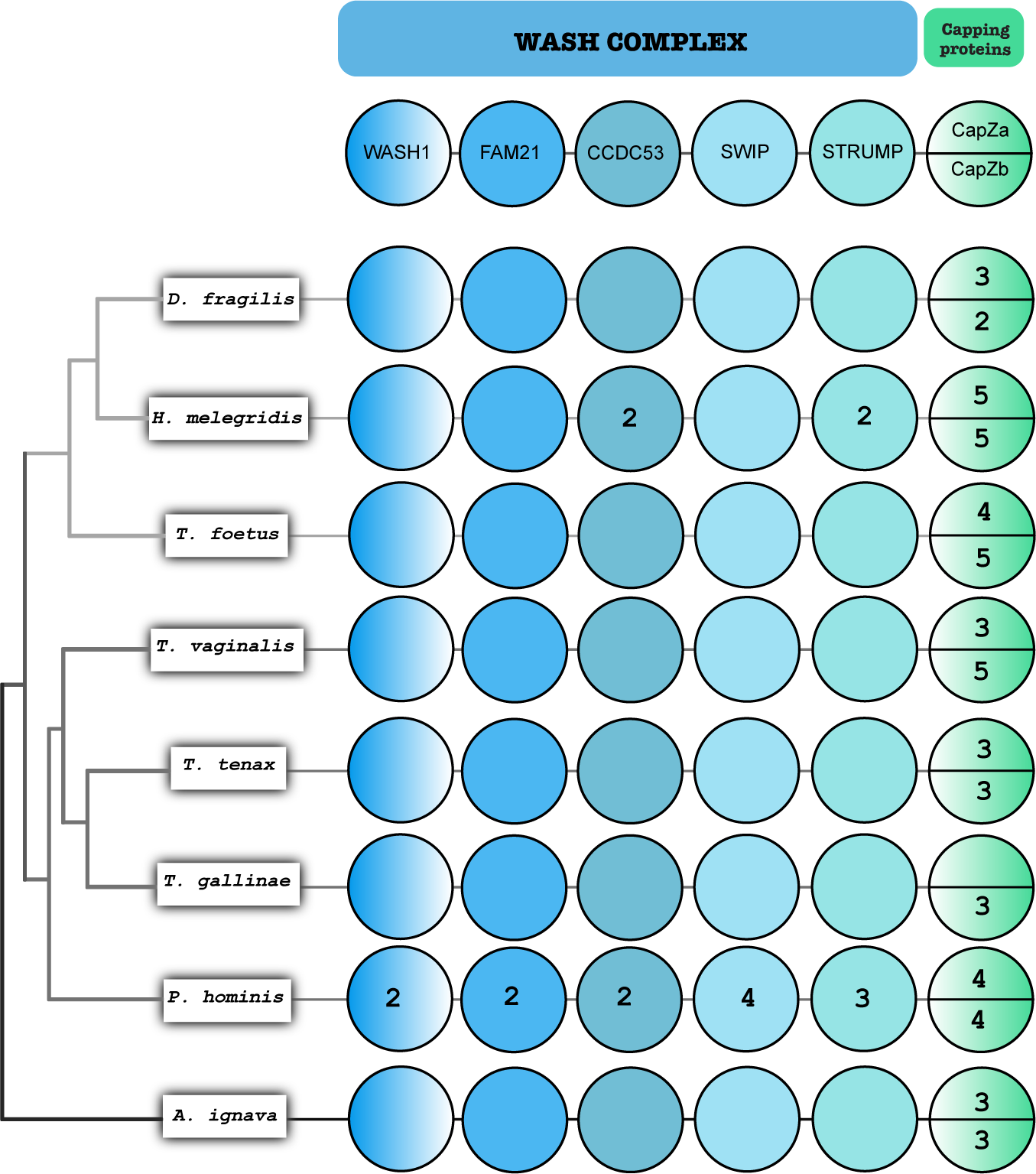
Coulson plot representing comparative genomics of the WASH complex and Capping proteins shared by Retromer and Retriever complexes in selected Parabasalid species and *A. ignava*. Areas filled with color demonstrate positive occurrence of the protein homologues, numbers indicate the paralogue count and colored sections with no numbers indicate a single paralogue. Each plot column is annotated with protein name at the top and the name of organism is mentioned on the left. WASH complex is colored in shades of blue and Capping proteins are colored in green. WASH subunits are labelled as follows; WASH1: WASHC1, FAM21: WASHC2, CCDC53: WASHC3, SWIP: WASHC4, STRUMP (strumpellin): WASHC5.

Based on our previous observation of the shared VPS29 protein being expanded like the Retromer complex in both Parabasalia and Anaeramoeba lineages (Figure 1A and Supporting Figure S1), we expected the WASH complex to showcase a similar pattern of evolution. However, to our surprise, we observed only a single gene homolog of WASH1 protein in nearly all the parabasalids and *A. ignava*. We observed the same pattern of evolution for FAM21 (WASHC2) as well, which is known to interact with VPS35 of the Retromer complex and CCDC93 of the Commander assembly with the Retriever complex (Jia et al., 2012), (Simonetti & Cullen, 2019). Similarly, CCDC53 (WASHC3) is also present as a single gene in five of the Parabasalia species and *A. ignava,* however, 2 paralogues were seen in *H. meleagridis* and *P. hominis* each. A single gene of SWIP (WASHC4) was seen in nearly all the parabasalids and *A. ignava*. SWIP was also recently identified to mediate the recruitment of the WASH complex to the endomembrane, independent of the Retromer complex (Dostál et al., 2023). The final WASH complex component: Strumpellin / STRUMP (WASHC5) also demonstrates a similar fashion of evolution, with single genes identified in the majority of the parabasalids and *A. ignava*. Like CCDC53, STRUMP is also present as two and three paralogues in *H. meleagridis* and *P. hominis* respectively. Hence, the WASH complex follows a similar pattern of evolution to the Retriever complex. We also confirmed species-specific duplication events specifically in *H. meleagridis* and *P. hominis*, by phylogenetic analysis of the WASH complex proteins in Parabasalia members (Supporting Figure S3).

Additionally, the dimeric Capping proteins; CapZα and CapZβ known to associate with the WASH complex and promote endosomal maturation (Wang et al., 2021) were also investigated. We observed an expansion of CapZα ranging from three genes in *D. fragilis, T. vaginalis, T. tenax,* and *A. ignava* to five genes in *H. meleagridis.* Four genes were seen in *T. foetus* and *P. hominis.* Only a single gene was identified in *T. gallinae*. For CapZβ, we observed an expansion ranging from two genes in *D. fragilis* to five genes in *H. meleagridis, T. foetus,* and *T. vaginalis*. An expansion of three genes was seen in *T. tenax, T. gallinae,* and *A. ignava* and expansion of four genes was seen in *P. hominis*.

Finally, we investigated the evolutionary pattern of the CCC complex (Figure 5) consisting of 13 proteins involving ten COMMD proteins, CCDC22, CCDC93 and DENND10 proteins exclusively interacting with only the Retriever complex and forming a megacomplex known as the Commander assembly. CCC complex plays a crucial role in the Retriever machinery by forming a link between the Retriever complex and the WASH complex to mediate actin polymerization (Simonetti & Cullen, 2019), (Healy et al., 2023).

**Figure 5:**
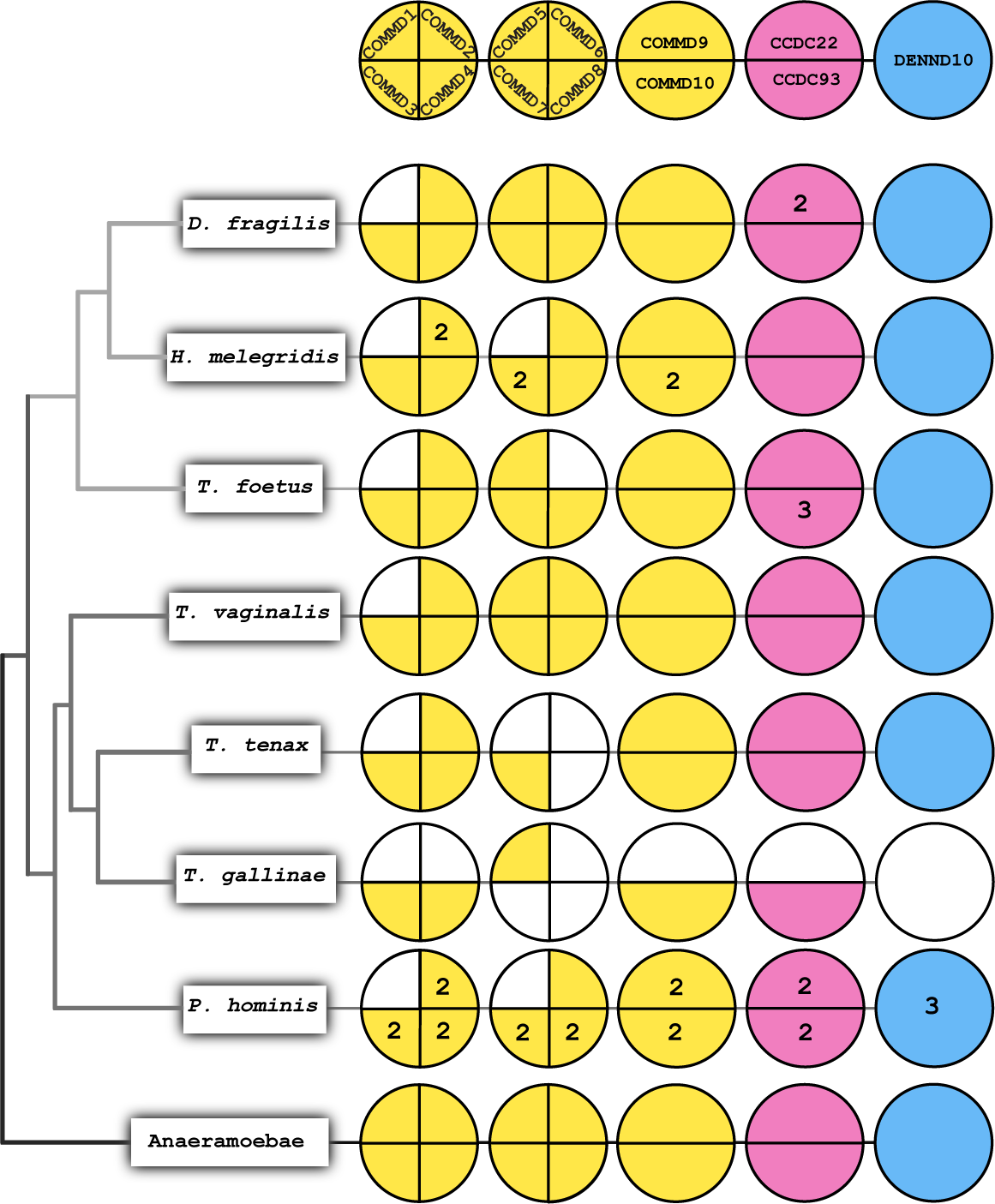
Coulson plot representing comparative genomics of the CCC sub-complex; together with Retriever complex known as the Commander complex. This complex consists of 13 proteins. Each plot column is annotated with protein name at the top and the name of organism is mentioned on the left. Areas filled with color demonstrate positive occurrence of the protein homologues, numbers indicate the paralogue count and colored sections with no numbers indicate a single paralogue. Sections left white or colorless indicate the absence of protein homologues. Corrective analysis for searches in *A. ignava*, with *A. flamelloides* is together mentioned as the Anaeramoeba lineage on the left in the list of all the assessed organisms.

We observed that the Anaeramoeba lineage shows the presence of all the CCC complex proteins, while COMMD1 of the CCC complex was seen to be completely lost in all the parasitic parabasalid members. We found single gene homologs of all the CCC complex proteins in the Anaeramoeba lineage (surveyed in *A. ignava* or *A. flamelloides*) and in nearly all the Parabasalids. However, we found an expansion of two genes for all the identified CCC complex proteins in *P. hominis* except for three paralogues identified for DENND10 and the presence of a single gene of COMMD4. A similar pattern of expansion was observed in *H. meleagridis* too, an expansion of two genes was seen for COMMD2, COMMD7, and COMMD10 and the rest of the CCC complex proteins identified are present as single homologs. Most of the expansions of COMMD proteins seen in *H. meleagridis* and *P. hominis*, like the WASH complex appear to be a result of species-specific duplication events (Supporting Figure S4).

In other parabasalids, the CCC complex proteins majorly contain single genes for each protein identified in our genomic survey while showing a complete loss of COMMD1. *T. tenax* appears to have lost COMMD5, COMMD6 and COMMD8. *T. gallinae* is seen to be missing COMMD1, 2, 6, 7, 8, and 9 along with CCDC22 and DENND10 which could have been a result of its poorer BUSCO score (Maciejowski et al., 2023). Among Tritrichomonadea, *T. foetus* appears to have lost COMMD4 along with COMMD1, while containing single homologs of other CCC complex proteins except for an expansion of three genes of CCDC93. *D. fragilis* is seen to have a single gene homolog for all the CCC complex proteins except for the expansion of two genes of CCDC22 and like other parabasalids to have lost the COMMD1 protein. A phylogenetic characterization of all the COMMD proteins identified in Anaeramoeba and Parabasalia members was conducted using human proteins as verified markers (Supporting Figure S4).

### 2.4 A possible answer to the mystery of the missing Sorting Nexins in Parabasalia lineage

Both the Retromer and Retriever complex machineries use Sorting Nexins to recruit the trimeric complexes at the endosomal membrane for its interaction with the endomembrane cargo proteins to be recycled, meanwhile also promoting the deformation of the membrane for vesicle formation (Gallon & Cullen, 2015). In mammals, SNX1,2,5 and 6 of the SNX-BAR category with the PX and BAR domain assist the Retromer complex (Collins et al., 2005), (Shi et al., 2006). Recently, SNX3 of the SNX-PX category was confirmed as a functional Sorting Nexin of the Retromer complex assembly with only a PX domain (Leneva et al., 2021). SNX27 and SNX17 are examples of SNX-FERM sorting nexins associated with Retromer assembly and Retriever assembly respectively (Chandra et al., 2021), (Healy et al., 2023), (Gallon et al., 2014). Thus, it was puzzling that no such SNX proteins were identified in prior analyses (Koumandou et al., 2011), (Gallon & Cullen, 2015). To investigate this mystery, we decided to explore this in all the parabasalids and *A. ignava* to find Sorting Nexins using Comparative genomics (Figure 1A).

We found that in *A. ignava*, the canonical Sorting Nexins; SNX-BAR are observed to have ten paralogues while having five paralogues of SNX-PX. We found SNX-FERM proteins to be completely absent in both the Parabasalia and Anaeramoeba lineages. In all the Parabasalia members, we observed 3-5 genes of SNX-PX proteins, however, SNX-BAR proteins were seen to be completely absent. Additionally, we also found 2-5 genes of ‘BAR only’ proteins in all the parabasalids, which are missing in the Anaeramoeba lineage. BAR-only proteins represent homology toward the C-terminus BAR domain of the fungal VPS5 (Figure 1B). Upon critically looking into the genome assembly of *T. vaginalis*, the SNX-PX homologs do not appear to be in the vicinity of BAR-only homologs and are also located on different chromosomes. This suggests that the SNX-PX and BAR-only proteins in parabasalids are independent proteins and possibly not a result of fragmentation in the genome. We also attempted phylogenetic analysis for the identified SNX proteins. Unfortunately, the tree showed no resolution (Data not shown). Thus, we have identified putative candidates for Sorting Nexins that could help mitigate the loss of standard eukaryotic Sorting Nexins in parasitic parabasalids.

### 2.5 Retromer and Retriever complexes localize differently from one another in the cell of *T. vaginalis*

The Retriever complex has only been characterized by molecular cell biological techniques in mammalian systems. Therefore, we decided to test whether these complexes act similarly in *T. vaginalis* as in mammals. We conducted sub-cellular localization of Retromer and Retriever complexes to distinguish the trafficking pathways using Immuno-fluorescence confocal microscopy (Figure 6). Since both the machineries had differential specificities to the endomembrane cargoes, we hypothesized them both to demonstrate differential localizations in *T. vaginalis,* with the Retromer complex recycling its cargo to the TGN and the plasma membrane and the Retriever complex recycling its cargo only to the plasma membrane.

**Figure 6:**
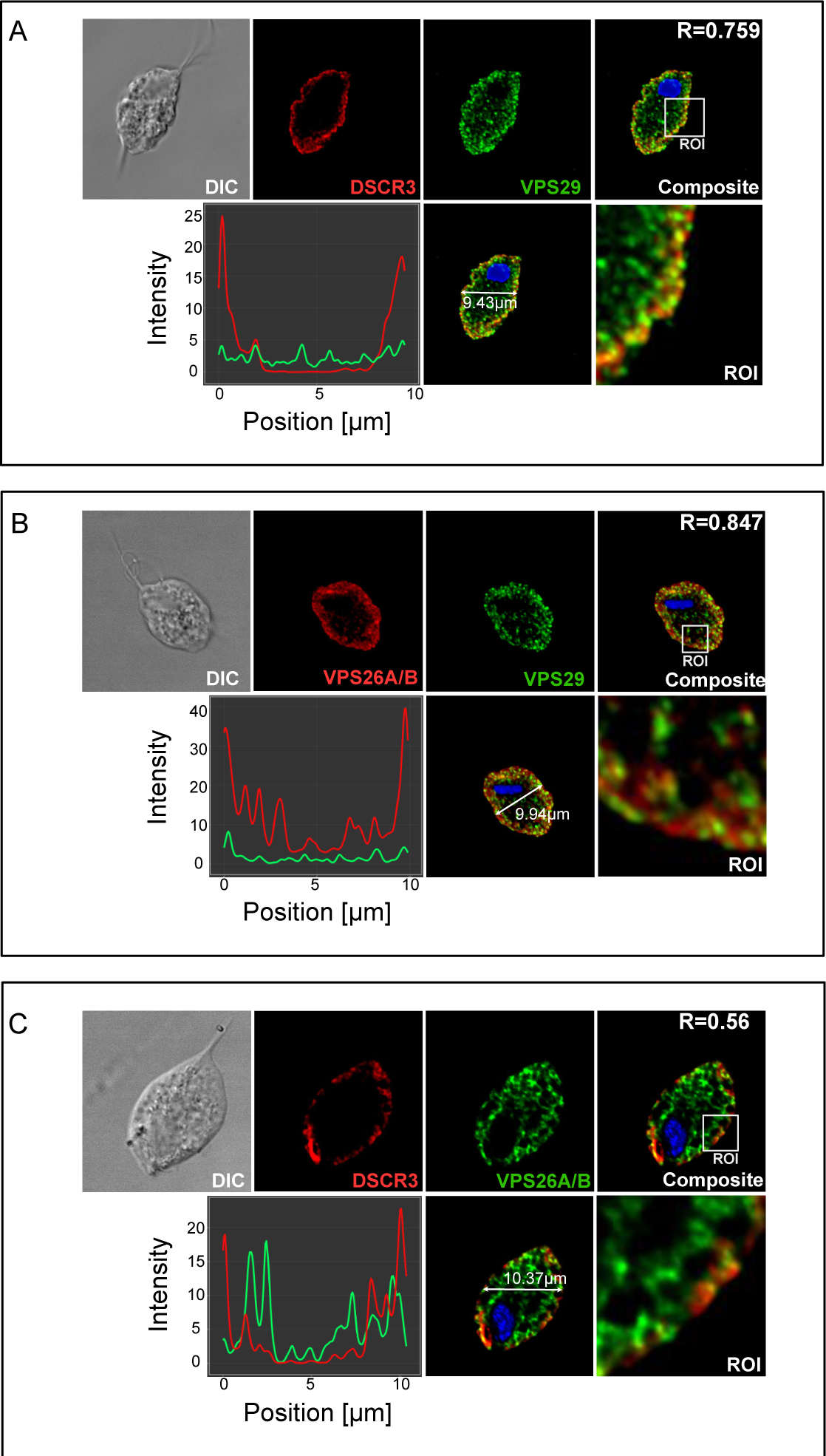
Differential cellular localization of Retromer and Retriever complex in *Trichomonas vaginalis*. **A)** Immuno-fluorescence co-localization of the Retriever specific DSCR3 with common subunit VPS29. Z-stack composite and channel Z-stacks of Immuno-fluorescence confocal microscopy images of *T. vaginalis* cell labelled with HAHA-DSCR3 (red) and V5-VPS29 (green). Cell position vs channel intensity plot, cell composite with scale and enlarged Region of Interest (ROI). **B)** Immuno-fluorescence co-localization of the Retromer specific VPS26A/B and shared subunit VPS29. Immuno-fluorescence confocal microscopy images of *T. vaginalis* cell labelled with HAHA-VPS26A/B (red) and V5-VPS29 (green). Cell position vs channel intensity plot, cell composite with scale and enlarged ROI. **C)** Immuno-fluorescence co-localization of the Retromer and Retriever complex. Immuno-fluorescence confocal microscopy images of *T. vaginalis* cell labelled with HAHA-DSCR3 (red) and V5-VPS6A/B (green). Cell position vs channel intensity plot, cell composite with scale and enlarged ROI.

**Figure 7:**
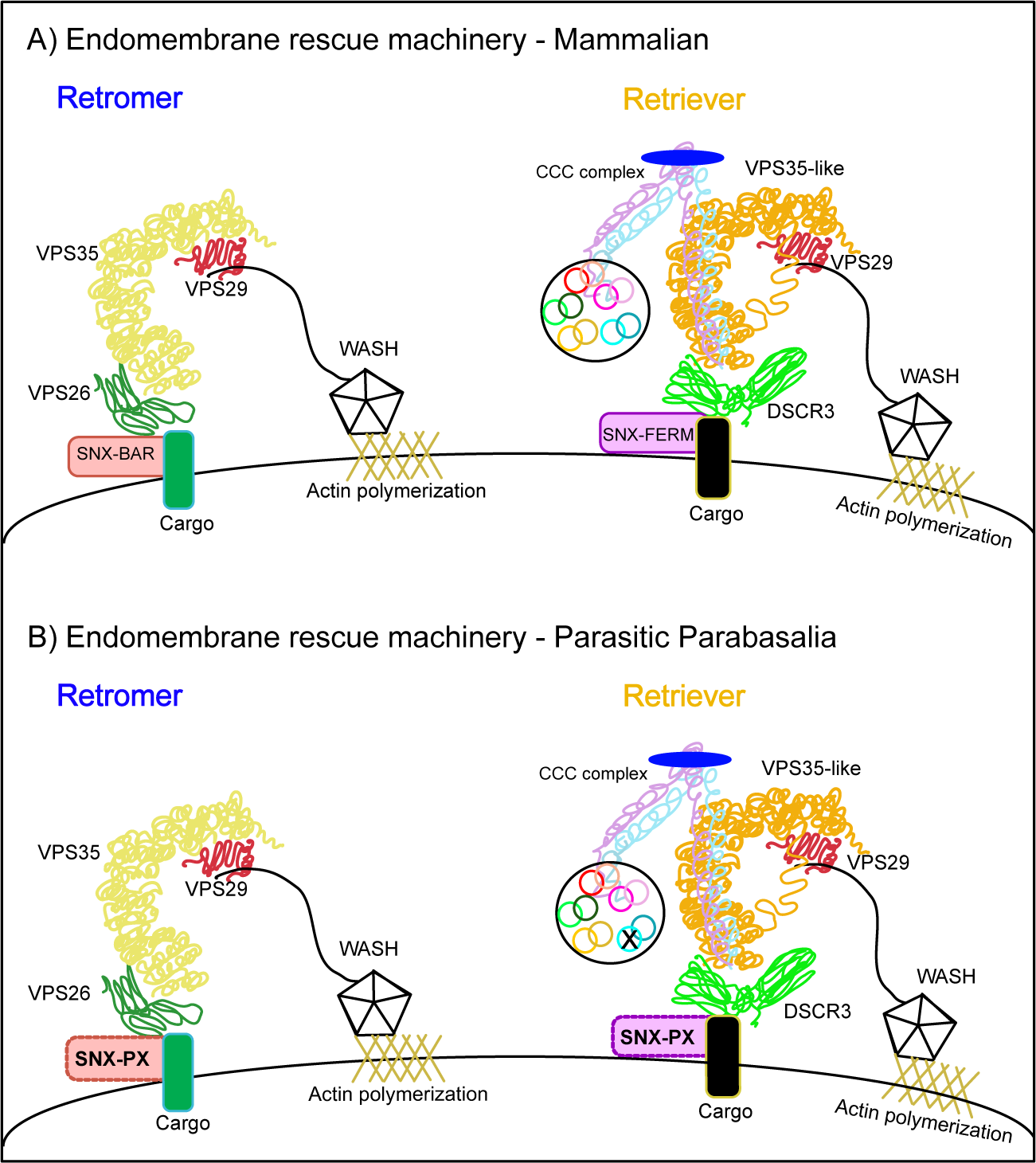
Sorting nexin hypothesis. **A)** Cartoon depiction of the canonical Retromer complex and Retriever complex machineries on the sorting endosomal membrane of a Mammalian system; ideally *Homo sapiens* Canonically, Sorting nexins in Retromer complex have additional BAR domain and the Sorting nexins for Retriever complex sorting have additional FERM domain, labelled as such. **B)** Cartoon depiction of reconstruction of Retromer and Retriever complex machineries in the parasitic Parabasalid system; ideally *Trichomonas vaginalis*. Missing COMMD1 of the CCC complex is marked with an X. Sorting nexins are proposed to be SNX-PX for both the complexes.

**Figure 8:**
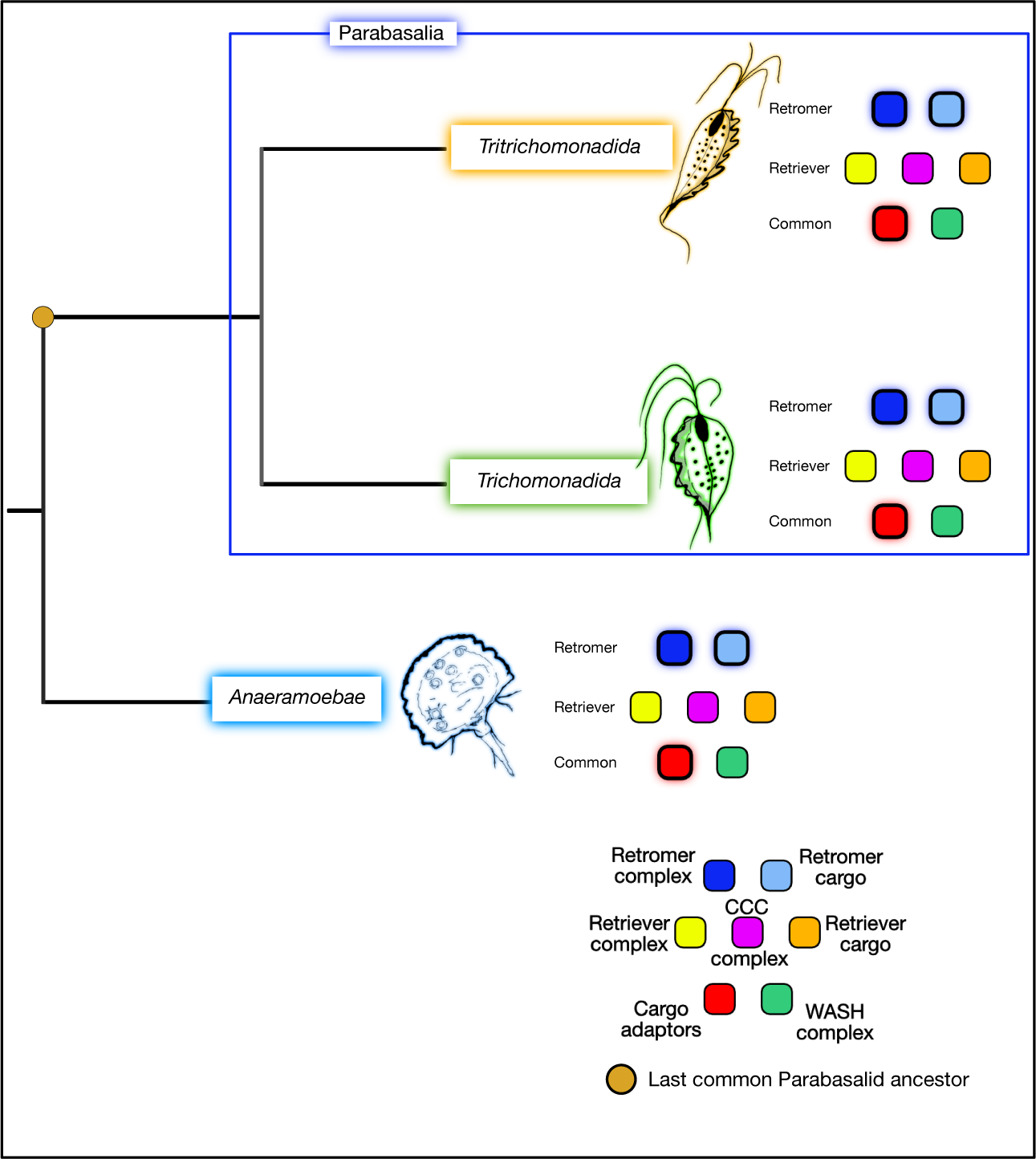
Conclusion of phylogenomics in Parabasalia and Anaeramoeba lineages; Outer glow of the coloured boxes represent gene expansion of the proteins belonging to specific complexes.

We co-expressed the Retriever complex specific protein; DSCR3 tagged with double human influenza hemagglutinin (HA) tag with the shared protein between Retromer and Retriever complexes; VPS29 tagged with V5 tag (Figure 6A). We observed DSCR3 to be distributed along the cell membrane (red channel). While VPS29 (green channel) co-localized significantly with DSCR3 on the cell surface, it was also observed in internal vesicles. The composite image confirms the co-localization of DSCR3 with VPS29 as can also be seen in the Region of Interest (ROI) image (Figure 6A-1). The represented image showed the co-localization parameter; Pearson co-efficient (R-value) of 0.759 for the complete cell.

Next, we studied the co-localization of VPS26A/B of the Retromer complex with the shared VPS29 protein. Like above, VPS29 was tagged with V5 while VPS26A/B was tagged with a double HA tag (Figure 6B). We observed VPS26A/B distributed along the cell membrane as well as inside the cells on the internal vesicles (red channel). Similarly, VPS29 (green channel) is also localized on the cell membrane and internal vesicles. From the composite image and the ROI, VPS26A/B and VPS29 co-localize internally on the membranes of vesicles, most arguably endosomes along with their distribution on the cell membrane (Figure 6B-1). The represented image showed an R-value of 0.874 for the complete cell.

Finally, we co-expressed HA-tagged VPS26A/B of the Retromer complex with the V5-tagged DSCR3 of the Retriever complex (Figure 6C). As expected, we observed DSCR3 (red channel) to be localized abundantly along the plasma membrane while VPS26A/B (green channel) was seen to be localized internally as well as towards the plasma membrane, both these localizations were consistent with Figures 6A & 6B. As can be seen from the Composite and ROI images, the Retromer and Retriever complex appears to co-localize only in certain spots on the plasma membrane (Figure 6C-1), significantly less than the membrane co-localization of DSCR3 with the shared VPS29 protein (Figure 6A). The represented image showed an R-value of 0.56 for the complete cell. We also provide intensity plots for each co-localization analysis determining the channel intensities in different cellular locations, clearly showing the Retromer complex being formed at the cell surface as well as internally meanwhile, the Retriever complex is formed only on the cell surface (Figure 6A/B/C-2).

## 3. DISCUSSION

Among the previous studies conducted on the Retromer complex, only one pan-eukaryotic comparative genomic survey has reported the presence of an endomembrane rescue machinery in Parabasalids showing a unique expansion of all the trimeric Retromer complex proteins in *T. vaginalis* whereas other Metamonada species with the presence of Retromer complex lack such an evolution (Koumandou et al., 2011). Despite its essential role in the membrane trafficking system in eukaryotes, more than a decade later there has still been no further investigation to study such a peculiar phenomenon in this significant human parasite. Our studies provide the first comprehensive investigation of the complete endomembrane rescue machinery in not only *T. vaginalis* but all the other parasitic parabasalids and their free-living sister of Anaeramoeba lineage.

Our investigations provide the first evidence for the presence of the Retriever complex machinery in Parabasalia and Anaeramoeba and we showed that its evolution is distinct from that of the Retromer complex. As the Retromer complex has been identified in *T. vaginalis*, we aimed to investigate if the homologous Retriever complex is also present in *T. vaginalis* and if it follows a similar pattern of expanded evolution. To increase the robustness, we conducted our genomic survey in all the available parasitic parabasalids and Anaeramoeba. To our surprise, DSCR3 and VPS35-L proteins of the Retriever complex did not appear to be expanded as the Retromer complex proteins; VPS26A/B and VPS35. This contrast in evolutionary pattern was found in *A. ignava* alongside all the parasitic parabasalids. The VPS29 protein shared between both these trimeric complexes is seen to be expanded like VPS26A/B and VPS35 of the Retromer complex in all the investigated taxa. Based on our *in-silico* investigations we speculate that the evolutionary pattern of the shared VPS29 protein is linked to the Retromer complex. This suggests that the evolution of the Retromer complex precedes the evolution of the Retriever complex, while DSCR3 and VPS35-L appeared as a result of ancestral duplication events in LECA and interacted with VPS29 to form a new endomembrane rescue machinery. In another instance, this is validated by the presence of VPS29 in all the Fungal species, while DSCR3 and VPS35-L are absent in the entire fungal lineage (McNally et al., 2017).

Based on our phylogenetic analyses to study the expansion of the Retromer complex (Supporting Figures S1 & S2), we also propose that due to a duplication event in LPCA, VPS29, and VPS35 evolved in Parabasalids with two orthologues each. Thus, giving all the parasitic Parabasalids, VPS29A/VPS29B and VPS35A/VPS35B. Meanwhile, ancestral duplication event in the evolution of VPS26A/B appears to have occurred only in Trichomonadea, but not in Tritrichomonadea of the Parabasalia lineage.

We show the presence of the Commander assembly consisting of the CCC complex along with the Retriever complex. The evolution of the CCC complex is as unexpanded as DSCR3 and VPS35-L, which portrays a synchronized evolution of both these interacting complexes in the Retriever-mediated rescue machinery. This is validated by the fact that the CCC complex is also absent in the fungal lineage and does not interact with the Retromer assembly. Hence, it is exclusive only to the Retriever complex machinery (McNally et al., 2017), (Healy et al., 2023). Another shared component between the Retromer and the Retriever complex that we subjected to genomic survey is the pentameric WASH complex, which is responsible for actin polymerization forming the endomembrane tubules and endomembrane maturation making it highly essential to the function of both the complexes. Due to its association with the Retriever as well as the Retromer complex, we expected its pattern of evolution to resonate with the shared VPS29 protein. However, to our surprise, we saw that the evolution of the WASH complex is like that of the Retriever complex in Parabasalia and Anaeramoeba lineages. It also critically differs from the pan-eukaryotic presence of the Retromer complex and demonstrates a distribution similar to the Retriever complex, while being absent in the fungal lineage (McNally et al., 2017). Based on our investigations we suggest that the distinct evolution trajectories of the Retromer complex and the Retriever complex could have driven the evolutionary patterns of these conserved accessory components. However, the difference in the evolution of shared components; VPS29 and the WASH complex remains to be assessed.

We also provide proof for the presence of putative Sorting Nexins in parabasalids and Anaeramoeba previously thought to be lost in *T. vaginalis*. Very recently, it was confirmed that SNX3, a Sorting Nexin containing only the PX domain is a functional cargo adaptor for the Retromer complex independently capable of Retromer-mediated cargo rescue (Cui et al., 2019), (Strochlic et al., 2007). Based on this information and our genomic survey, we hypothesize that the putative SNX-PX homologs identified in Parabasalids are highly divergent Sorting Nexins acting as the functional cargo adaptors for both the Retromer complex and the Retriever complex (Figure 1). We hope that further studies and deeper investigations will evaluate and validate our hypothesis.

The Retromer complex and the Retriever complex demonstrate different sub-cellular localizations in the model parasitic Parabasalid; *T. vaginalis*. Having confirmed the distinct evolutionary patterns for the Retromer and the Retriever complex machineries, we evaluated if they differed in their rescue pathways in *T. vaginalis* (Figure 6). Based on our observations, we suggest that the Retromer complex rescues its specific endomembrane cargoes either back to the cell surface or internally to the trans-Golgi network, while the Retriever complex rescues its endomembrane cargoes primarily to the cell surface. This confirms that both these complexes demonstrate functional homology in this parasitic protist lineage. This is the first report of a cellular localization study of the Retromer complex in any metamonad species and the first non-metazoan report for cellular localization of the Retriever complex.

In conclusion, we provide a comprehensive investigation of the endomembrane rescue machinery in parasitic parabasalids and their free-living sister; Anaeramoeba. It reveals the distinct evolution of the Retriever complex from the Retromer complex, with shared VPS29 protein exhibiting an expanded evolutionary pattern like the Retromer complex and the shared WASH complex and the Retriever-specific CCC complex exhibiting an evolutionary pattern like the Retriever complex. Additionally, the presence of putative Sorting nexins in Parabasalids and Anaeramoeba challenges previous beliefs. The study also demonstrates different sub-cellular localizations of the Retromer and Retriever complex. These findings contribute to our understanding of the evolutionary dynamics and functional conservation of both the rescue machineries in parasitic protists, and we hope that further investigations in other parasitic systems will evaluate and validate our hypothesis.

## 4. Experiment procedures

### 4.1 Taxa genomic and transcriptome dataset acquisition

Genomes, protein datasets, and Transcriptomes were acquired from National Centre for Biotechnology Information (NCBI) for *T. vaginalis* (Jane M. Carlton & Robert P. Hirt, n.d.), *T. foetus* (Benchimol et al., 2017), *H. meleagridis* (Palmieri et al., 2021), *D. fragilis*, *P. hominis*, *T. tenax,* and *T. gallinae* were obtained from NCBI (Handrich et al., 2019). *Anaeramoeba ignava* and *Anaeramoeba flamelloides* transcriptome data were obtained from FigShare (Stairs et al., 2021). The list and sources of the datasets are provided in the Supporting table S1.

### 4.2 Homology searches and domain analysis

Homology searches of the components in Endomembrane Retrograde transport machineries and their cargoes were conducted using **A**nalysis of **Mo**lecular **E**volution with **BA**tch **E**ntry (AMOEBAE) (Barlow et al., 2023). *Homo sapiens* and *Saccharomyces cerevisiae* protein sequences from NCBI were used as queries for the initial homology searches. Forward searching was done with BLASTp and tBLASTn, e-value max limit parameter was set to 0.05 to sort all the positive hits. AMOEBAE then confirmed the positive hits by performing reciprocal BLAST searches with the same e-value cut-off. Positive hits from this were used to prepare Hidden Markov Models (HMM) to conduct HMMER searches using AMOEBAE for all the queries across above obtained Parabasalia and Anaeramoebae genome and transcriptomes datasets.

The final positive hits were validated and confirmed for homology using HHPred (Zimmermann et al., 2018) and InterProScan (Paysan-Lafosse et al., 2023) for the domain presence. The confirmed positive hits with accession numbers of the proteins are listed in Supporting table S1.

### 4.3 Phylogenetic analyses and tree construction

All the protein sequence alignments were created using MAFFT v7.505 (Katoh et al., 2019). For visualization of the alignments, a fast and lightweight software; AliView was used (Larsson, 2014). The alignments were trimmed to obtain the informative regions using Block Mapping and Gathering with entropy (BMGE) (Criscuolo & Gribaldo, 2010). Partial sequences identified were ignored from the final analyses. All the alignments are available upon request. Phylogenetic analyses were carried out by maximum likelihood approach using IQTree2 (Minh et al., 2020). IQTree Model finder was used to choose the best-fit models for all the analyses (Kalyaanamoorthy et al., 2017). Ultrafast bootstrapping (-B -N 1000) and non-parametric bootstrapping (-b -N 1000) support values were generated with IQTree2. FigTree *v1.4.4* (http://tree.bio.ed.ac.uk/software/figtree/) was used to visualize and analyze the phylogenetic data.

### 4.4 Cell culture and cultivation

*Trichomonas vaginalis* strain Tv17-48 was used for this study (Kulda et al., 1982). The cells were axenically cultured at 37 °C in TYM (<u>T</u>ryptone <u>Y</u>east extract <u>M</u>altose) medium at pH 6.2 with 10% heat inactivated horse serum. For the growth and selection of co-transfected cells, the TYM media was supplemented with 100 mg/ml of Geneticin and 40 mg/ml of Puromycin.

### 4.5 Expression vector construction and nucleofection into the *T. vaginalis*

VPS26A/B (TrichDB: TVAG_160820, NCBI: XP_001324499.1) and DSCR3 (TrichDB: TVAG_332210, NCBI: XP_001299315.1) were expressed in *T. vaginalis* with C-terminal hemagglutinin (HA) tag using vector pTagVag-HA-Neo (Hrdy et al., 2004). The genes were expressed under the control of their respective native promoters of 250 bp upstream of the coding sequence. VPS26A/B and VPS29 (TrichDB: TVAG_378930, NCBI: XP_001583400.1) were co-expressed with C-terminal V5 tag in double transfectants using vector pTagVag-V5-Pur (Štácová et al., 2018). VPS29 expression was under the control of the native promoter of 300bp upstream. The genes were amplified by PCR using genomic DNA of Tv17-48 as a template and cloned in the vectors via SacII and BamHI restriction sites. The vectors were co-transfected into the Tv17-48 trichomonad cells by nucleofection using Lonza’s Human T cell Nucleofector Kit according to (Zimmann et al., 2022), (Janssen et al., 2018). The gene sequence and PCR primers are provided in Supporting table S1.

### 4.6 Immunofluorescence Microscopy and Image Analysis

*Trichomonas vaginalis* transfected cells were harvested by centrifugation at 1,200 x g for 10 minutes. The cells were then washed with Phosphate buffered solution (PBS) and fixed using 2% formaldehyde as described by (Woessner & Dawson, 2012). The fixed cells attached to Poly L-lysine treated coverslips were incubated overnight at 4 °C with primary antibodies; rabbit polyclonal anti-HA antibody (Santa-Cruz Biotechnology Inc, catalogue: SC-805) and mouse monoclonal anti-V-5 antibody (Thermo Fisher Scientific, catalogue: 37-7500). Alexa Fluor 594 donkey anti-rabbit (Life Technologies) and Alexa Fluor 488 donkey anti-mouse (Life Technologies) were used as secondary antibodies. The nucleus was stained with 4′,6-diamidino-2-phenylindole (DAPI). Slides were observed using a Leica TCS SP8 confocal laser scanning microscope with the 63x oil immersion objective. Captured images were deconvolved using Huygens Professional v19.10 (https://svi.nl/Huygens-Professional) and processed using ImageJ/Fiji open-source software (Schindelin et al., 2012). Co-localization analysis was performed using a voxel-based co-localization analysis tool of Huygens Professional v19.10 in Z-stack to calculate Pearson’s correlation coefficient (PCC/R) using Costes method.

## 5. Data availability

All the data is provided in the figures, figure legends and Supporting information.

## 6. Supporting information

This article contains supporting information.

## 7. Acknowledgements

We thank Michaela Marcinčiková for her technical assistance. The authors acknowledge the Imaging Methods Core Facility at BIOCEV, Charles University, Czech Republic, supported by the Ministry of Education, Youth and Sport of the Czech Republic (LM2023050 Czech-BioImaging) for their support and assistance with Confocal microscopy.

## 8. Funding and additional information

The research was funded by the Charles University to APS (GAUK 354622), JK (UNCE/SCI/12, UNCE24/SCI/011), and JT (Cooperatio Biology), and European Regional Development Fund and Ministry of Education, Youth and Sports of the Czech Republic (No. CZ.02.1.01/0.0/0.0/16_019/0000759). Research in the Dacks Lab was supported by grants from the Natural Sciences and Engineering Research Council of Canada (RES0043758, and RES0046091).

## 9. Author contributions

APS, JBD and JT designed the studies. APS performed Comparative genomics, Phylogenetics and data illustration. APS and JK performed microscopy. APS, JBD and JT carried out data analyses and wrote the manuscript. All the authors read and approved the final manuscript.

## 12. Abbreviations

VPS: (Vacuolar protein sorting-associated protein)
DSCR3: (Down syndrome critical region protein 3)
CIMPR: (Cation independent Mannose-6-phosphate receptor)
DPAP: (Dipeptidyl aminopeptidase)
KEX2: (Kexin 2)
LDLR: (Low-density lipoprotein receptor)
LRP: (Low-density lipoprotein receptor-related protein)
SNX: (Sorting Nexins)
PX: (phox homology)
BAR: (Bin, Amphiphysin, and Rvs)
FERM: (4.1/ezrin/radixin/moesin)
WASH: (<u>W</u>iskott-<u>A</u>ldrich syndrome protein and <u>S</u>CAR <u>H</u>omologue)
CCDC: (Coiled-coil domain-containing protein)
SWIP: (Strumpellin and WASH-interacting protein)
STRUMP: (Strumpellin)
CapZα/β: (F-actin capping protein subunit alpha/beta)
COMMD: (<u>Co</u>pper <u>M</u>etabolism <u>M</u>URR1 [Mouse U2af1-rs1 region 1] <u>D</u>omain)
DENND10: (<u>D</u>ifferentially <u>E</u>xpressed in <u>N</u>ormal and <u>N</u>eoplastic cells-containing <u>D</u>omain protein 10)
TGN: (trans-Golgi network)
MVB: (multi-vesicular bodies)
ER: (Endoplasmic reticulum).

---

Supporting figure S1: Maximum likelihood phylogenetic analysis tree was constructed using IQTree (Best fit: Q.yeast+G4) with 24 sequences and 180 sites for VPS29 expansion events in parasitic Parabasalids. This phylogenetic tree includes all the orthologues identified from homology searches in Parabasalids and *A. ignava*. The clades in green represents orthologues of VPS29 identified in the parasitic Parabasalia lineage. The clade in pink represents paralogues identified in *A. ignava* where the tree is rooted. A partial sequence of VPS29 from *A. ignava* was eliminated from this analysis for the robustness of the dataset. The red dot represents an ancestral duplication event during the speciation of the parasitic Parabasalids and the blue dots represent species-specific duplication events. Support values for each node are presented in the format of UFB/NP. Branches with support values of 100 for both ultrafast and NP bootstrapping are represented with black solid triangles.

Supporting figure S2: Maximum likelihood phylogenetic analysis tree was constructed using IQTree (Best fit: Q.yeast+I+G4) with 21 sequences and 545 sites for VPS35 expansion events in parasitic Parabasalids. This phylogenetic tree includes all the orthologues identified from homology searches in Parabasalia and *A. ignava.* This tree was rooted at mid-point. The clades in blue represents orthologues of VPS35 identified in the parasitic Parabasalia lineage. The clade in pink represents paralogues identified in *A. ignava*. Two partial sequences of VPS35 from *A. ignava* were eliminated from this analysis to maintain the robustness of the dataset. The red dot represents an ancestral duplication event during the speciation of the parasitic Parabasalids and the blue dots represent species specific duplication events. Support values for each node are presented in the format of UFB/NP. Branches with support values of 100 for both ultrafast and NP bootstrapping are represented with black solid triangles.

Supporting figure S3: Maximum likelihood phylogenetic analysis tree was constructed using IQTree (Best fit: LG+F+G4) with 44 sequences and 424 sites for the WASH complex in the parasitic Parabasalids and *A. ignava*. The phylogenetic tree was constructed using all the WASH complex protein homologues and orthologues identified from Comparative genomic analysis (Figure 4). WASH complex protein sequences from *H. sapiens* were used as reference for the phylogenetic characterization of each WASH complex protein. The tree was rooted at WASH1. Support values for each node are depicted in the format UFB/NP. Branches with support values of 100 for both ultrafast and NP bootstrapping are represented with black solid triangles.

Supporting figure S4: Maximum likelihood phylogenetic analysis tree was constructed using IQTree (Best fit: LG+F+G4) with 89 sequences and 137 sites for characterization of COMMD1-10 proteins of the CCC complex in Parabasalia and Anaeramoeba lineages. The phylogenetic tree was constructed using all the COMMD protein homologues and orthologues identified from Comparative genomic analysis (Figure 5). COMMD protein sequences from *H. sapiens* were used as reference for the phylogenetic characterization of each COMMD protein. The tree was rooted at COMMD1 clade (represented in red) and COMMD2-10 clades are represented in yellow. Support values for each node are depicted in the format UFB/NP. Branches with support values of 100 for both ultrafast and NP bootstrapping are represented with black solid triangles.

## References

Adl, S. M., Bass, D., Lane, C. E., Lukeš, J., Schoch, C. L., Smirnov, A., Agatha, S., Berney, C., Brown, M. W., Burki, F., Cárdenas, P., Čepička, I., Chistyakova, L., Campo, J., Dunthorn, M., Edvardsen, B., Eglit, Y., Guillou, L., Hampl, V., … Zhang, Q. (2019). Revisions to the Classification, Nomenclature, and Diversity of Eukaryotes. Journal of Eukaryotic Microbiology, 66(1), 4–119. 10.1111/jeu.12691

Arighi, C. N., Harmell, L. M., Aguilar, R. C., HaP, C. R., & Bonifacino, J. S. (2004). Role of the mammalian retromer in sorting of the cation-independent mannose 6-phosphate receptor. Journal of Cell Biology, 165(1), 123–133. 10.1083/jcb.200312055

Barlow, L. D., Maciejowski, W., More, K., Terry, K., Vargová, R., Záhonová, K., & Dacks, J. B. (2023). Compara-ve Genomics for Evolu-onary Cell Biology Using AMOEBAE: Understanding the Golgi and Beyond (pp. 431–452). 10.1007/978-1-0716-2639-9_26

Benchimol, M., de Almeida, L. G. P., Vasconcelos, A. T., de Andrade Rosa, I., Reis Bogo, M., Kist, L. W., & de Souza, W. (2017). DraP Genome Sequence of Tritrichomonas foetus Strain K. Genome Announcements, 5(16). 10.1128/genomeA.00195-17

Benchimol, M., Diniz, J. A. P., & Ribeiro, K. (2000). The fine structure of the axostyle and its associations with organelles in Trichomonads. Tissue and Cell, 32(2), 178–187. 10.1054/nce.2000.0102

Burki, F., Roger, A. J., Brown, M. W., & Simpson, A. G. B. (2020). The New Tree of Eukaryotes. Trends in Ecology & Evolution, 35(1), 43–55. 10.1016/j.tree.2019.08.008

Céza, V., Kotyk, M., Kubánková, A., Yubuki, N., Šťáhlavský, F., Silberman, J. D., & Čepička, I. (2022). Free-living Trichomonads are Unexpectedly Diverse. Protist, 173(4), 125883. 10.1016/j.protis.2022.125883

Chandra, M., Kendall, A. K., & Jackson, L. P. (2021). Toward Understanding the Molecular Role of SNX27/Retromer in Human Health and Disease. In Frontiers in Cell and Developmental Biology (Vol. 9). Frontiers Media S.A. 10.3389/fcell.2021.642378

Chen, K.-E., Healy, M. D., & Collins, B. M. (2019). Towards a molecular understanding of endosomal trafficking by Retromer and Retriever. 10.1111/tra.12649/Abstract

Collins, B. M., Skinner, C. F., Watson, P. J., Seaman, M. N. J., & Owen, D. J. (2005). Vps29 has a phosphoesterase fold that acts as a protein interaction scaffold for retromer assembly. Nature Structural & Molecular Biology, 12(7), 594–602. 10.1038/nsmb954

Criscuolo, A., & Gribaldo, S. (2010). BMGE (Block Mapping and Gathering with Entropy): a new soPware for selection of phylogenetic informative regions from multiple sequence alignments. BMC Evolutionary Biology, 10(1), 210. 10.1186/1471-2148-10-210

Cui, Y., Carosi, J. M., Yang, Z., Ario, N., Kerr, M. C., Parton, R. G., Sargeant, T. J., & Teasdale, R. D. (2019). Retromer has a selective function in cargo sorting via endosome transport carriers. Journal of Cell Biology, 218(2), 615–631. 10.1083/jcb.201806153

Derivery, E., Sousa, C., Gautier, J. J., Lombard, B., Loew, D., & Gautreau, A. (2009). The Arp2/3 Activator WASH Controls the Fission of Endosomes through a Large Multiprotein Complex. Developmental Cell, 17(5), 712–723. 10.1016/j.devcel.2009.09.010

Dostál, V., Humhalová, T., Beránková, P., Pácalt, O., & Libusová, L. (2023). <scp>SWIP</scp> mediates retromer-independent membrane recruitment of the <scp>WASH</scp> complex. Traffic, 24(5), 216–230. 10.1111/tra.12884

Edgar, A. J., & Polak, J. M. (2000). Human Homologues of Yeast Vacuolar Protein Sorting 29 and 35. Biochemical and Biophysical Research Communications, 277(3), 622–630. 10.1006/bbrc.2000.3727

Edwards, T., Burke, P., Smalley, H., & Hobbs, G. (2016). Trichomonas vaginalis: Clinical relevance, pathogenicity and diagnosis. Critical Reviews in Microbiology, 42(3), 406–417. 10.3109/1040841X.2014.958050

Gallon, M., Clairfeuille, T., Steinberg, F., Mas, C., Ghai, R., Sessions, R. B., Teasdale, R. D., Collins, B. M., & Cullen, P. J. (2014). A unique PDZ domain and arrestin-like fold interaction reveals mechanistic details of endocytic recycling by SNX27-retromer. Proceedings of the National Academy of Sciences, 111(35). 10.1073/pnas.1410552111

Gallon, M., & Cullen, P. J. (2015). Retromer and sorting nexins in endosomal sorting. Biochemical Society Transac-ons, 43(1), 33–47. 10.1042/BST20140290

HaP, C. R., Sierra, M. de la L., Bafford, R., Lesniak, M. A., Barr, V. A., & Taylor, S. I. (2000). Human Orthologs of Yeast Vacuolar Protein Sorting Proteins Vps26, 29, and 35: Assembly into Multimeric Complexes. Molecular Biology of the Cell, 11(12), 4105–4116. 10.1091/mbc.11.12.4105

Handrich, M. R., Garg, S. G., Sommerville, E. W., Hirt, R. P., & Gould, S. B. (2019). Characterization of the BspA and Pmp protein family of trichomonads. Parasites & Vectors, 12(1), 406. 10.1186/s13071-019-3660-z

Harrison, M. S., Hung, C.-S., Liu, T., Christiano, R., Walther, T. C., & Burd, C. G. (2014). A mechanism for retromer endosomal coat complex assembly with cargo. Proceedings of the National Academy of Sciences of the United States of America, 111(1), 267–272. 10.1073/pnas.1316482111

Healy, M. D., McNally, K. E., Butkovič, R., Chilton, M., Kato, K., Sacharz, J., McConville, C., Moody, E. R. R., Shaw, S., Planelles-Herrero, V. J., Yadav, S. K. N., Ross, J., Borucu, U., Palmer, C. S., Chen, K.-E., Croll, T. I., Hall, R. J., Caruana, N. J., Ghai, R., … Cullen, P. J. (2023). Structure of the endosomal Commander complex linked to Ritscher-Schinzel syndrome. Cell, 186(10), 2219–2237.e29. 10.1016/j.cell.2023.04.003

Hrdy, I., Hirt, R. P., Dolezal, P., Bardonová, L., Foster, P. G., Tachezy, J., & Martin Embley, T. (2004). Trichomonas hydrogenosomes contain the NADH dehydrogenase module of mitochondrial complex I. Nature, 432(7017), 618–622. 10.1038/nature03149

Huotari, J., & Helenius, A. (2011). Endosome maturation. The EMBO Journal, 30(17), 3481– 3500. 10.1038/emboj.2011.286

Jane M. Carlton, & Robert P. Hirt. (n.d.). DraV Genome Sequence of the Sexually TransmiYed Pathogen Trichomonas vaginalis.

Janssen, B. D., Chen, Y.-P., Molgora, B. M., Wang, S. E., Simoes-Barbosa, A., & Johnson, P. J. (2018). CRISPR/Cas9-mediated gene modification and gene knock out in the human-infective parasite Trichomonas vaginalis. Scientific Reports, 8(1), 270. 10.1038/s41598-017-18442-3

Jia, D., Gomez, T. S., Billadeau, D. D., & Rosen, M. K. (2012). Multiple repeat elements within the FAM21 tail link the WASH actin regulatory complex to the retromer. Molecular Biology of the Cell, 23(12), 2352–2361. 10.1091/mbc.e11-12-1059

Kalyaanamoorthy, S., Minh, B. Q., Wong, T. K. F., von Haeseler, A., & Jermiin, L. S. (2017). ModelFinder: fast model selection for accurate phylogenetic estimates. Nature Methods, 14(6), 587–589. 10.1038/nmeth.4285

Kang, S., Tice, A. K., Stairs, C. W., Jones, R. E., Lahr, D. J. G., & Brown, M. W. (2021). The integrin-mediated adhesive complex in the ancestor of animals, fungi, and amoebae. Current Biology, 31(14), 3073–3085.e3. 10.1016/j.cub.2021.04.076

Katoh, K., Rozewicki, J., & Yamada, K. D. (2019). MAFFT online service: multiple sequence alignment, interactive sequence choice and visualization. Briefings in Bioinforma-cs, 20(4), 1160–1166. 10.1093/bib/bbx108

Kerr, M. C., Bennetts, J. S., Simpson, F., Thomas, E. C., Flegg, C., Gleeson, P. A., Wicking, C., & Teasdale, R. D. (2005). A Novel Mammalian Retromer Component, Vps26B. Traffic, 6(11), 991–1001. 10.1111/j.1600-0854.2005.00328.x

Koumandou, V. L., Klute, M. J., Herman, E. K., Nunez-Miguel, R., Dacks, J. B., & Field, M. C. (2011). Evolutionary reconstruction of the retromer complex and its function in Trypanosoma brucei. Journal of Cell Science, 124(9), 1496–1509. 10.1242/jcs.081596

Krieger, J. N. (1995). Trichomoniasis in men: old issues and new data. Sexually TransmiYed Diseases, 22(2), 83–96.

Kulda, J., Vojtechovska, M., Tachezy, J., Demes, P., & Kunzova, E. (1982). Metronidazole resistance of Trichomonas vaginalis as a cause of treatment failure in trichomoniasis--A case report. Sexually TransmiYed Infec-ons, 58(6), 394–399. 10.1136/sn.58.6.394

Larsson, A. (2014). AliView: a fast and lightweight alignment viewer and editor for large datasets. Bioinformacs, 30(22), 3276–3278. 10.1093/bioinformatics/btu531

Leneva, N., Kovtun, O., Morado, D. R. G. Briggs, J. A., & Owen, D. J. (2021). STRUCTURAL BIOLOGY Architecture and mechanism of metazoan retromer:SNX3 tubular coat assembly. In Sci. Adv (Vol. 7). https://www.science.org

Lucas, M., Gershlick, D. C., Vidaurrazaga, A., Rojas, A. L., Bonifacino, J. S., & Hierro, A. (2016). Structural Mechanism for Cargo Recognition by the Retromer Complex. Cell, 167(6), 1623–1635.e14. 10.1016/j.cell.2016.10.056

Maciejowski, W. J., Gile, G. H., Jerlström-Hultqvist, J., & Dacks, J. B. (2023). Ancient and pervasive expansion of adaptin-related vesicle coat machinery across Parabasalia. International Journal for Parasitology, 53(4), 233–245. 10.1016/j.ijpara.2023.01.002

Malik, S.-B., Brochu, C. D., Bilic, I., Yuan, J., Hess, M., Logsdon, J. M., & Carlton, J. M. (2011). Phylogeny of parasitic parabasalia and free-living relatives inferred from conventional markers vs. Rpb1, a single-copy gene. PloS One, 6(6), e20774. 10.1371/journal.pone.0020774

Mallam, A. L., & Marcotte, E. M. (2017). Systems-wide Studies Uncover Commander, a Multiprotein Complex Essential to Human Development. Cell Systems, 4(5), 483–494. 10.1016/j.cels.2017.04.006

Mann, J. R., McDermott, S., Barnes, T. L., Hardin, J., Bao, H., & Zhou, L. (2009). Trichomoniasis in Pregnancy and Mental Retardation in Children. Annals of Epidemiology, 19(12), 891–899. 10.1016/j.annepidem.2009.08.004

McClelland, R. S., Sangaré, L., Hassan, W. M., Lavreys, L., Mandaliya, K., Kiarie, J., Ndinya-Achola, J., Jaoko, W., & Baeten, J. M. (2007). Infection with *Trichomonas vaginalis* Increases the Risk of HIV-1 Acquisition. The Journal of Infectious Diseases, 195(5), 698– 702. 10.1086/511278

McNally, K. E., & Cullen, P. J. (2018). Endosomal Retrieval of Cargo: Retromer Is Not Alone. In Trends in Cell Biology (Vol. 28, Issue 10, pp. 807–822). Elsevier Ltd. 10.1016/j.tcb.2018.06.005

McNally, K. E., Faulkner, R., Steinberg, F., Gallon, M., Ghai, R., Pim, D., Langton, P., Pearson, N., Danson, C. M., Nägele, H., Morris, L. L., Singla, A., Overlee, B. L., Heesom, K. J., Sessions, R., Banks, L., Collins, B. M., Berger, I., Billadeau, D. D., … Cullen, P. J. (2017). Retriever is a multiprotein complex for retromer-independent endosomal cargo recycling. Nature Cell Biology, 19(10), 1214–1225. 10.1038/ncb3610

Minh, B. Q., Schmidt, H. A., Chernomor, O., Schrempf, D., Woodhams, M. D., von Haeseler, A., & Lanfear, R. (2020). IQ-TREE 2: New Models and Efficient Methods for Phylogenetic Inference in the Genomic Era. Molecular Biology and Evolution, 37(5), 1530–1534. 10.1093/molbev/msaa015

Nothwehr, S. F., & Hindes, A. E. (1997). The yeast *VPS5/GRD2* gene encodes a sorting nexin-1-like protein required for localizing membrane proteins to the late Golgi. Journal of Cell Science, 110(9), 1063–1072. 10.1242/jcs.110.9.1063

Palmieri, N., de Jesus Ramires, M., Hess, M., & Bilic, I. (2021). Complete genomes of the eukaryotic poultry parasite Histomonas meleagridis: linking sequence analysis with virulence / attenuation. BMC Genomics, 22(1), 753. 10.1186/s12864-021-08059-2

Paysan-Lafosse, T., Blum, M., Chuguransky, S., Grego, T., Pinto, B. L., Salazar, G. A., Bileschi, M. L., Bork, P., Bridge, A., Colwell, L., Gough, J., HaP, D. H., Letunić, I., Marchler-Bauer, A., Mi, H., Natale, D. A., Orengo, C. A., Pandurangan, A. P., Rivoire, C., … Bateman, A. (2023). InterPro in 2022. Nucleic Acids Research, 51(D1), D418–D427. 10.1093/nar/gkac993

Pipaliya, S. V., Santos, R., Salas-Leiva, D., Balmer, E. A., Wirdnam, C. D., Roger, A. J., Hehl, A. B., Faso, C., & Dacks, J. B. (2021). Unexpected organellar locations of ESCRT machinery in Giardia intestinalis and complex evolutionary dynamics spanning the transition to parasitism in the lineage Fornicata. BMC Biology, 19(1), 167. 10.1186/s12915-021-01077-2

Raiborg, C., & Stenmark, H. (2009). The ESCRT machinery in endosomal sorting of ubiquitylated membrane proteins. Nature, 458(7237), 445–452. 10.1038/nature07961

Schindelin, J., Arganda-Carreras, I., Frise, E., Kaynig, V., Longair, M., Pietzsch, T., Preibisch, S., Rueden, C., Saalfeld, S., Schmid, B., Tinevez, J.-Y., White, D. J., Hartenstein, V., Eliceiri, K., Tomancak, P., & Cardona, A. (2012). Fiji: an open-source plaÄorm for biological-image analysis. Nature Methods, 9(7), 676–682. 10.1038/nmeth.2019

Seaman, M. N. J. (2021). The Retromer Complex: From Genesis to Revelations. In Trends in Biochemical Sciences (Vol. 46, Issue 7, pp. 608–620). Elsevier Ltd. 10.1016/j.nbs.2020.12.009

Seaman, M. N. J., Michael McCaffery, J., & Emr, S. D. (1998). A Membrane Coat Complex Essential for Endosome-to-Golgi Retrograde Transport in Yeast. The Journal of Cell Biology, 142(3), 665–681. 10.1083/jcb.142.3.665

Shi, H., Rojas, R., Bonifacino, J. S., & Hurley, J. H. (2006). The retromer subunit Vps26 has an arrestin fold and binds Vps35 through its C-terminal domain. Nature Structural & Molecular Biology, 13(6), 540–548. 10.1038/nsmb1103

Simone, B., & Cullen, P. J. (2019). Actin-dependent endosomal receptor recycling. In Current Opinion in Cell Biology (Vol. 56, pp. 22–33). Elsevier Ltd. 10.1016/j.ceb.2018.08.006

Štácová, J., Rada, P., Meloni, D., Žárský, V., Smutná, T., Zimmann, N., Harant, K., Pompach, P., Hrdý, I., & Tachezy, J. (2018). Dynamic secretome of Trichomonas vaginalis: Case study of β-amylases. Molecular & Cellular Proteomics, 17(2), 304–320. 10.1074/mcp.RA117.000434

Stairs, C. W., Táborský, P., Salomaki, E. D., Kolisko, M., Pánek, T., Eme, L., Hradilová, M., Vlček, Č., Jerlström-Hultqvist, J., Roger, A. J., & Čepička, I. (2021). Anaeramoebae are a divergent lineage of eukaryotes that shed light on the transition from anaerobic mitochondria to hydrogenosomes. Current Biology, 31(24), 5605–5612.e5. 10.1016/j.cub.2021.10.010

Steinberg, F., Gallon, M., Winfield, M., Thomas, E. C., Bell, A. J., Heesom, K. J., Tavaré, J. M., & Cullen, P. J. (2013). A global analysis of SNX27-retromer assembly and cargo specificity reveals a function in glucose and metal ion transport. Nature Cell Biology, 15(5), 461–471. 10.1038/ncb2721

Strochlic, T. I., Setty, T. G., Sitaram, A., & Burd, C. G. (2007). Grd19/Snx3p functions as a cargo-specific adapter for retromer-dependent endocytic recycling. The Journal of Cell Biology, 177(1), 115–125. 10.1083/jcb.200609161

Sutcliffe, S., Giovannucci, E., Alderete, J. F., Chang, T. H., Gaydos, C. A., Zenilman, J. M., De Marzo, A. M., Willett, W. C., & Platz, E. A. (2006). Plasma antibodies against Trichomonas vaginalis and subsequent risk of prostate cancer. Cancer Epidemiology Biomarkers and Prevention, 15(5), 939–945. 10.1158/1055-9965.EPI-05-0781

Teasdale, R. D., & Collins, B. M. (2012). Insights into the PX (phox-homology) domain and SNX (sorting nexin) protein families: structures, functions and roles in disease. Biochemical Journal, 441(1), 39–59. 10.1042/BJ20111226

Wang, D., Ye, Z., Wei, W., Yu, J., Huang, L., Zhang, H., & Yue, J. (2021). Capping protein regulates endosomal trafficking by controlling F-actin density around endocytic vesicles and recruiting RAB5 effectors. ELife, 10. 10.7554/eLife.65910

Woessner, D. J., & Dawson, S. C. (2012). The Giardia Median Body Protein Is a Ventral Disc Protein That Is Critical for Maintaining a Domed Disc Conformation during Attachment. Eukaryo-c Cell, 11(3), 292–301. 10.1128/EC.05262-11

Zimmann, N., Rada, P., Žárský, V., Smutná, T., Záhonová, K., Dacks, J., Harant, K., Hrdý, I., & Tachezy, J. (2022). Proteomic Analysis of Trichomonas vaginalis Phagolysosome, Lysosomal Targeting, and Unconventional Secretion of Cysteine Peptidases. Molecular & Cellular Proteomics, 21(1), 100174. 10.1016/j.mcpro.2021.100174

Zimmermann, L., Stephens, A., Nam, S.-Z., Rau, D., Kübler, J., Lozajic, M., Gabler, F., Söding, J., Lupas, A. N., & Alva, V. (2018). A Completely Reimplemented MPI Bioinformatics Toolkit with a New HHpred Server at its Core. Journal of Molecular Biology, 430(15), 2237– 2243. 10.1016/j.jmb.2017.12.007

